# TLR4 regulation in human fetal membranes as an explicative mechanism of a pathological preterm case

**DOI:** 10.1101/2021.06.28.450131

**Authors:** Corinne Belville, Flora Ponelle-Chachuat, Marion Rouzaire, Christelle Gross, Bruno Pereira, Denis Gallot, Vincent Sapin, Loïc Blanchon

**Affiliations:** Team “Translational approach to epithelial injury and repair,” GReD, Université Clermont Auvergne, UMR6293 CNRS-U1103 INSERM, F-63000 Clermont-Ferrand, France; CHU Clermont-Ferrand, Biostatistics unit (DRCI) Department, F-63000 Clermont-Ferrand, France; CHU Clermont-Ferrand, Obstetrics and Gynecology Department, F-63000 Clermont-Ferrand, France; CHU Clermont-Ferrand, Biochemistry and Molecular Genetic Department, F-63000 Clermont-Ferrand, France

**Author notes:** Corresponding author: Dr Loic Blanchon, UFR de Médecine et des Professions Paramédicales, Bâtiment CRBC, GReD, TSA 50400, 28 place Henri-Dunant, F63001 Clermont-Ferrand Cedex 1, France.

**Keywords:** fetal membranes rupture, methylome, miRNA, transcriptome, TLR4

## Abstract

The integrity of human fetal membranes is crucial for harmonious fetal development throughout pregnancy. Their premature rupture is often the consequence of a physiological phenomenon previously exacerbated. Beyond all biological processes implied, inflammation is of primary importance and is qualified as “sterile” at the end of pregnancy. Complementary methylomic and transcriptomic strategies on amnion and choriodecidua explants taken from the altered (cervix zone) and intact fetal membranes at term and before labor were used in this study. By cross-analyzing genome-wide studies strengthened by *in vitro* experiments, we deciphered how the expression of Toll-like receptor 4 (TLR4), a well-known actor of pathological fetal membrane rupture, is controlled. Indeed, it is differentially regulated in the altered zone and between both layers by a dual mechanism: 1) the methylation of TLR4 and miRNA promoters and 2) targeting by miRNA (let-7a-2 and miR-125b-1) acting on the 3’-UTR of TLR4. Consequently, this study demonstrates that a fine regulation of TLR4 is required for sterile inflammation establishment at the end of pregnancy and that it may be dysregulated in the pathological premature rupture of membranes.

## INTRODUCTION

Placenta and fetal membranes are extra-embryonic tissues originally formed by trophoblastic cell differentiation. Although these organs are transitory, their integrities throughout pregnancy are essential for harmonious *in-utero* fetal development. Protecting the fetus (considered a semi-allogeneic graft), fetal membranes are composed of amnion and chorion, creating a 0.5 mm-thick layer surrounding the amniotic cavity. They are important for the protection of the fetus against infections ascending the genital tract and for the initiation of programmed term rupture (Joyce *et al*, 2009; Kendal-Wright, 2007). The site of rupture is a particular and unique zone of altered morphology (ZAM) situated around the cervix (McLaren *et al*, 1999; McParland *et al*, 2003), which along with a zone of intact morphology (ZIM) presents specific histological characteristics involving a substantial reduction of both layer thickness and cellular apoptosis (El Khwad *et al*, 2006) decreased the adherence of the amnion and the choriodecidua; the appearance of a swollen, spongy layer between them (Mauri *et al*, 2013) and epithelial to mesenchymal transition (EMT) (Silva *et al*, 2020). The evident global regional disorganization is principally a consequence of neutrophil and macrophage migrations (Osman *et al*, 2006) and collagen remodeling, which are essentially linked to an increased activity of matrix metalloproteinases (MMP) (McLaren *et al*, 2000). Furthermore, the release of pro-inflammatory cytokines due to innate immunity activates global inflammatory pathways, usually referred to as “sterile inflammation” (Gomez-Lopez *et al*, 2014). A sequential scenario occurs with the first chorion’s breaking, amnion invagination, and rupture (Gillaux *et al*, 2011).

Dysregulation of fetal membrane’s integrity or a premature activation of inflammatory pathways could lead to preterm premature rupture of membranes (PPROM). This is associated with a high mortality rate and significant morbidity in newborn survivors due to fetal prematurity and maternal complications (Goldenberg *et al*, 2008). Such pathology, affecting the amnion and chorion before 37 weeks of gestation, accounts for approximatively 30% of premature deliveries and 1–3 % of all births (Waters & Mercer, 2011). Deregulated genes (e.g., MMP1, ITGA11, and THBS2; (Wang *et al*, 2008; Yoo *et al*, 2018) miRNA, (Enquobahrie *et al*, 2016), and long chain non-coding RNA (lncRNA) (Luo *et al*, 2013)) expressions were demonstrated to correlate with PPROM appearance by affecting different biological pathways. Several causes have been found to increase the incidence of this pathology, where microorganisms ascending and invading the intra-amniotic cavity appear to be one of the most important explanations (Konwar *et al*, 2018; Musilova *et al*, 2015; Romero *et al*, 2006a). This leads to chorioamnionitis, triggering an early and acute inflammatory response and implying the involvement of intracellular or surface-expressed pattern recognition receptors (PRRs) as innate components of the immune system, including the most frequently described toll-like receptor (TLR) family (Kawai & Akira, 2010; Newton & Dixit, 2012).

To better understand the physiological and pathological rupture of membranes, the molecular study of global changes in gene expression can be accomplished using high-scale technics analyses. Only 12 % of the studies concerning gestational tissues have used fetal membranes. Furthermore, only 5 % investigated healthy pregnancies, whereas a physiological case could undoubtedly serve as an image of a premature exacerbated phenomenon in a pathological case. Ninety percent were conducted on pathological pregnancies, with 31 % specifically involving PPROM (Eidem *et al*, 2015). In term delivery, several researchers have observed an acute inflammatory signature in fetal membranes under different conditions: labor versus no labor (Haddad *et al*, 2006) in the placental amnion versus the reflected amnion with labor (Han *et al*, 2008) and in the choriodecidua (Stephen *et al*, 2015). Using human primary amnion mesenchymal cells treated with IL-1β, an elegant model of sterile inflammation (rapidly-activated by NF-κB responsive genes) was found to be associated with an inflammatory cascade (Li *et al*, 2011; Lim *et al*, 2012). Thus, in the ZAM, the sterile inflammation and/or extracellular matrix (ECM) remodeling and disorganization absolutely depends on global gene expression profiles in the amnion and the chorion. Surprisingly, only one unique transcriptome study focused on the ZIM or ZAM regions and amnion and chorion tissues, characterizing a specific differential expression in a spontaneous rupture at term with labor. Differences were observed in the chorion, though not in the amnion, specifically involving biological processes, such as extracellular matrix-receptor interaction and inflammation (Nhan-Chang *et al*, 2010). In addition to classical direct gene regulation by transcription factors, DNA methylation is another well-known epigenetic mechanism that can interfere with the transcriptional regulation of all RNA types (Kim *et al*, 2009). Concerning a physiological rupture, the differential methylation study regarding fetal membranes only focused on the amnion. Genome-wide methylation differed between the labor and non-labor groups at different pregnancy terms, particularly affecting the genes involved in cytokine production and gated channel activity, among others (Kim *et al*, 2013).

Because little is known about biochemical changes at term without labor (not influenced by mechanical stress) with regard to physiological conditions at the site of rupture (ZAM) and away from the site (ZIM), the purpose of our study was to exhaustively combine and correlate methylomic and transcriptomic analyses. We hypothesized that by cross-analyzing our genome-wide studies, we could explain the different levels of gene expression by comparing the amnion/choriodecidua and the ZIM/ZAM. After classifying genes into specific biological processes, we focused our study on inflammation. Under conditions at term without labor, we demonstrated that toll-like receptor 4 (TLR4), classically involved in the recognition of E.coli and triggering an inflammatory response in chorioamnionitis leading to PPROM (Medzhitov *et al*, 1997; Poltorak *et al*, 1998), was overexpressed in the ZAM choriodecidua compared to the ZAM amnion. Furthermore, we discovered that TLR4 regulation leads to layer and zone specificity. The latter occurred due to the hypomethylation of the TLR4 gene body in the ZAM choriodecidua, whereas its weak expression in the ZAM amnion layer was a direct consequence of the action of two hypomethylated miRNAs targeting the 3-UTR of TLR4: let-7a-2 and miR-125b-1. Therefore, the physiological choriodecidual over-expression of TLR4 could be exacerbated in PPROM, leading to the enhancement of the first step of an early scenario of a fetal membrane rupture.

## RESULTS

### The methylomic analysis of fetal membranes allows for defining the gene ontology classification for the ZAM zone

After identifying the differentially hypermethylated genes between the amnion and the choriodecidua on the whole genome between the ZIM and ZAM (Figure 1A), a biological process analysis was performed and represented by a four-way Venn diagram (Figure 1B). If the hypermethylated genes were clearly lower in the ZIM (1,746 genes) than the ZAM (9,830 genes), the specific genes only found in the ZIM (hyper- or hypomethylated; total number 98 [i.e., 10 + 88, illustrated by black circles]) did not exhibit any statistically significant result (p-value < 0.01 with a Bonferroni correction). In contrast, the specific genes modified in terms of methylation for the ZAM (total number 5,750 [i.e., 3,379 + 2,371, illustrated by white circles]) led to the discovery of the numerous biological processes detailed in Figure 1C. Precisely, the number of characterizations of an enrichment in GO term IDs were clearly more concentrated and were thus more biologically significant in the case where a definite gene was more methylated in the amnion than in the choriodecidua (mA > mC). The biological process term IDs containing the most important gene number were the G-protein coupled receptor signaling pathways (such as for defense response, detection of stimulus, or inflammatory response), which could be linked to sterile inflammation and to the onset of parturition related to fetal membrane ruptures.

**Figure 1:**
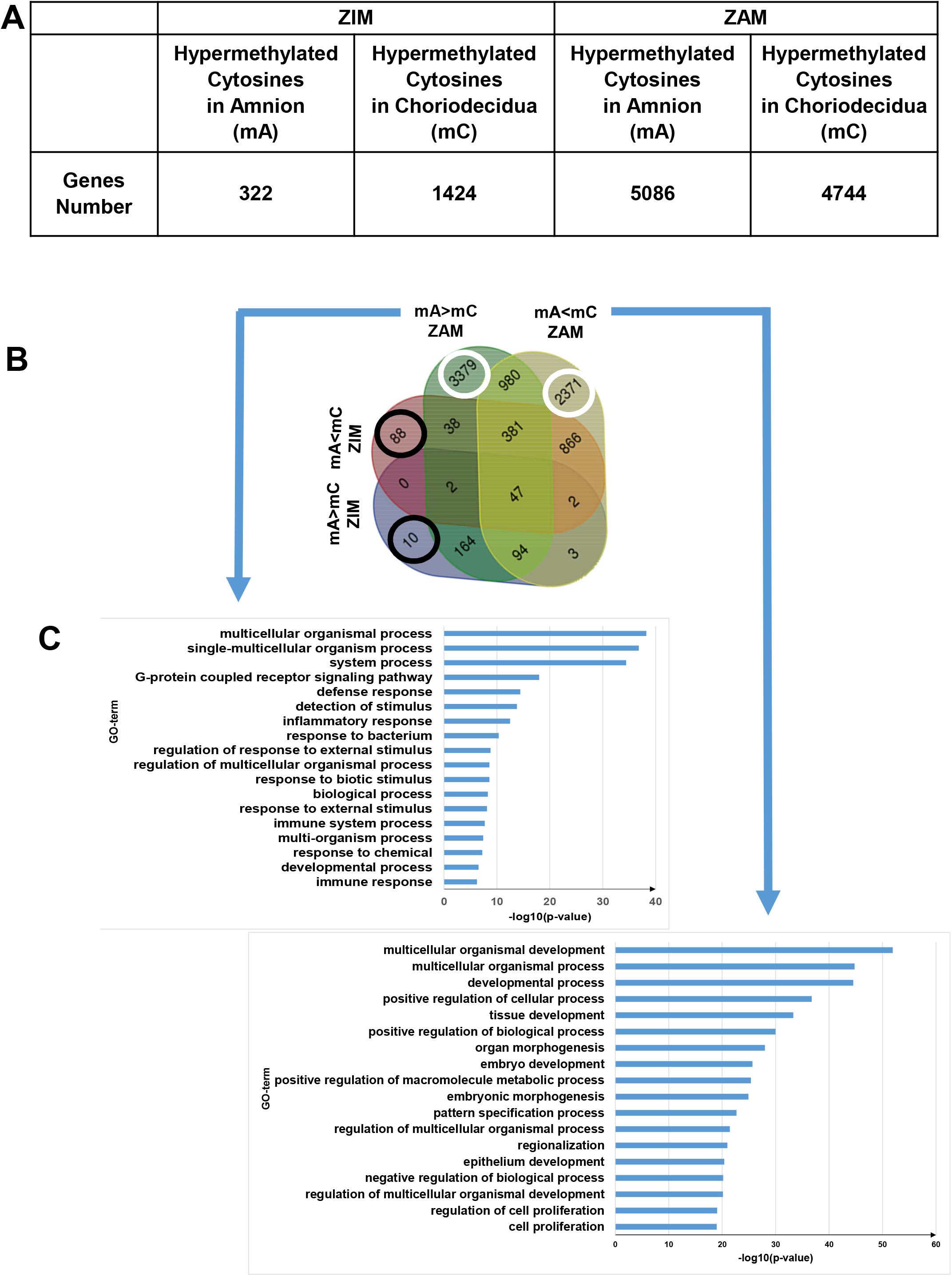
Differential cytosine methylation is analyzed for the ZIM and the ZAM. A Genes affected by differential methylation between the amnion and the choriodecidua, separately studied for the ZIM (left) and the ZAM (right). B Four-way Venn diagram representing the number of genes with hypermethylated cytosines in the ZIM and the ZAM according to the methylomic analyses. mA > mC: a specific gene was more methylated in the amnion than in the choriodecidua. mA < mC: a specific gene was less methylated in the amnion than in the choriodecidua. C GO-term classifications for genes observed specifically in the ZAM: mA > mC (left panel) and mA < mC (right panel). A Bonferroni correction was conducted for p-values < 0.01.

### The transcriptomic analysis of fetal membranes in the ZAM zone is more relevant for downregulated genes in the amnion compared to the choriodecidua

To supplement the results obtained from the methylome analysis, a transcriptomic study using the same samples was performed for the ZAM to compare the difference in gene expression between the amnion and the choridecidua. We observed that 501 and 145 genes specific to the ZAM were respectively down- and upregulated when the expression levels were compared between the amnion and the choriodecidua (a log2 fold change [FC] cut-off lower or higher than 2.8 [Figure 2, middle panel, supplementary Table S3]).

**Figure 2:**
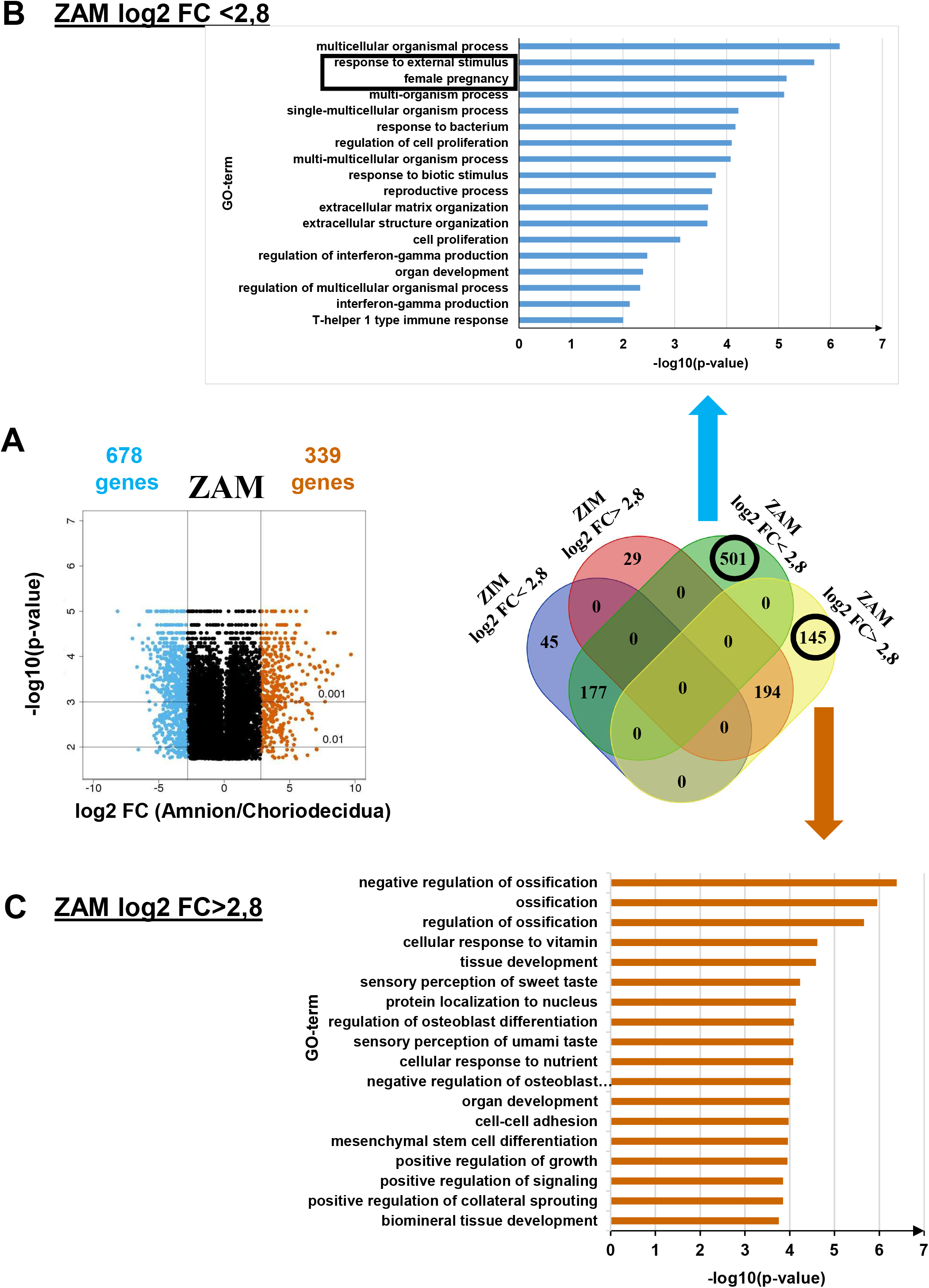
Transcriptomic assay is analyzed for the ZAM in fetal membranes. A Volcano plots represent the log10 adjusted p-values versus the log2 fold change (FC). Up- and downregulated genes are shown in red and blue, respectively, limited by│log_2_ FC│=2.8. They are classified in a four-way-Venn diagram representing the gene numbers in the ZIM and ZAM analyses with│log_2_ FC│=2.8. B GO-term classifications are shown for genes expressed only in the ZAM for log2 FC < 2.8 (Bonferroni correction for p-values < 0.01) C GO-term classifications are shown for genes expressed only in the ZAM for log2 FC > 2.8 (uncorrected p-value < 0.01).

The processes seemed to be more relevant and specific when the genes less expressed in the amnion than in the choriodecidua (i.e., log2 FC < 2.8) were considered. Indeed, genes could be grouped into two relevant GO terms: *response to external stimulus and female pregnancy* (Figure 2, top panel). In the case where genes were more expressed in the amnion than in the choriodecidua (i.e., log2 FC > 2.8), the biological processes exhibited no statistically significant result (p-value < 0.01 with a Bonferroni correction). The analyses undertaken with uncorrected p-values did not provide relevant information because conventional tissue development or ossification were observed (Figure 2, bottom panel).

### Combination of transcriptomic and methylomic results in the ZAM zone demonstrate that genes more expressed in the choriodecidua are linked to pregnancy pathologies

By cross-analyzing the results obtained from the two preceding analyses (methylome and transcriptome), a total of 26 genes were hypermethylated in the choriodecidua, and more were expressed in the amnion (supplementary Table S4). The biological process (GO term) analysis was not significant, which is likely due to the small number of studied genes and the fact that the underlying generic processes were linked only to urogenital abnormalities and not to pathological pregnancy, as confirmed by the MeSH disease terms (supplementary Table S5).

Conversely, 105 genes were hypermethylated in the amnion and were more expressed in the choriodecidua (Figure 3A and supplementary Table S6). They could be classified in MeSH disease terms linked directly to pregnancy pathologies, such as placenta diseases (trophoblastic neoplasms, pre-eclampsia, fetal growth retardation, or placenta accreta), female urogenital diseases, and pregnancy complications (supplementary Table S7). The GO term analysis clearly identified three complementary pathways: response to external stimulus, detection of LPSs, and inflammatory response. Interestingly, fetal membranes were sensitive to an external stimulus, such as Gram-negative bacterial molecules and LPSs, in relation to the inflammatory response. Of all the genes classified in these processes, TLR4 was the only one represented in all these biological processes and therefore seems to play a central role in parturition at term. To validate our *in-silico* observations and to pave the way for describing TLR4’s importance, immunofluorescence experiments were first conducted to confirm such a protein’s presence in the amnion and the choriodecidua of the ZAM (Figure 3B). Moreover, we established the ability of the choriodecidua and the amnion to respond to TLR4 activation after the binding of its natural ligand (LPS) by secreting IL-6 and TNF-α proteins in the culture medium (Figure 3C). Considering the important role of TLR4 in sterile or pathological inflammation (chorioamnionitis), the remainder of the study focused on the characterization and regulation of TLR4 expression in fetal membranes, as ascertained by the cross-analysis of the methylomic and transcriptomic results.

**Figure 3:**
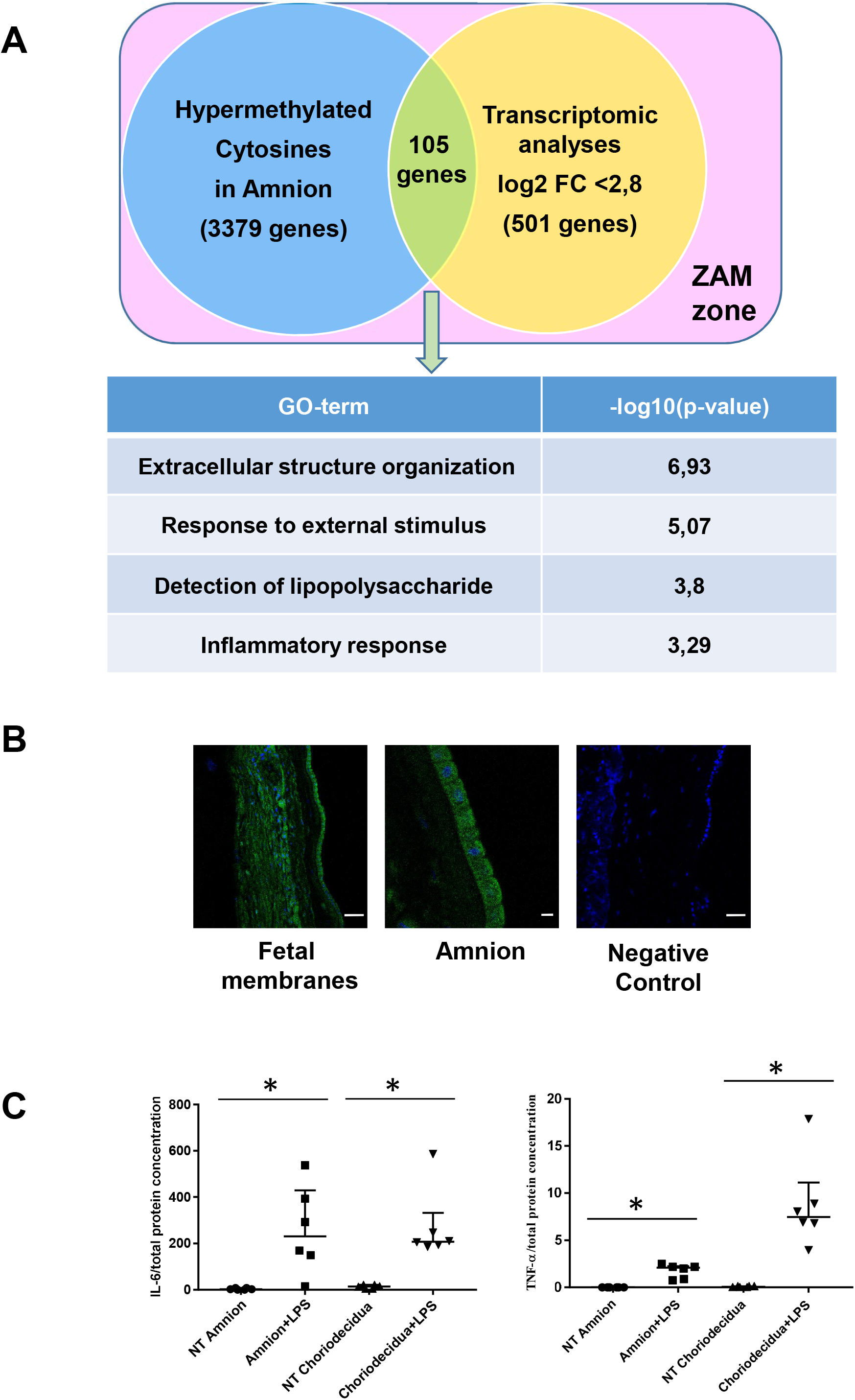
Common genes are observed between the mA > mC methylomic results and the choriodecidua/amnion transcriptomic analysis in the ZAM. A GO terms’ representative distribution and their log10 p-values are shown for the 105 common genes across both genome-wide studies. B Representative TLR4 immunofluorescence (green staining) in the ZAM of fetal membranes in the confocal analyses. Cell nuclei were visualized with Hoescht (blue) staining. A negative control was used without a primary antibody. Slides were observed at x250 magnification for total fetal membranes (left and right): scale bar: 50µm and x400 for the amnion (middle): scale bar: 20µm. C Luminex technology was used to detect physiological IL-6 and TNF-α levels and to demonstrate no (or a weak) presence of this interleukin in the amnion or the choriodecidua (NT condition). After treatment for 24 h with LPS (0.5µg/ml), the levels of IL-6 (n=6) and TNF-α (n=6) significantly increased. Median ± interquartile ranges are represented (Wilcoxon matched-pairs signed rank test, * p-value < 0.05).

### The tissue specificity of TLR4 regulation at the end of pregnancy in the weak ZAM is due to hypermethylation in the amnion

Focusing on the TLR4 genomic zone, five cytosines distributed on the promoter, body, and UTR were studied using a methylomic array. The statistical study of the cytosine methylation β-values of the nine genomic DNA samples demonstrated that the five were hypermethylated in the amnion compared to the choriodecidua (Figure 4A). These results were confirmed by the bisulphite treatment and enzymatic digestion (taking advantage of the Taq I restriction site present only in this CpG zone) of the amnion and choriodecidua samples with a focus on only one cytosine: cg 05429895 (Figure 4B). Furthermore, of the nine RNA samples, the link between methylation and expression revealed by the methylomic and trancriptomic arrays was confirmed by qRT-PCR (Figure 4C) and Western blot (Figure 4D) analyses, showing a higher expression of TLR4 in the choriodecidua than in the amnion.

**Figure 4:**
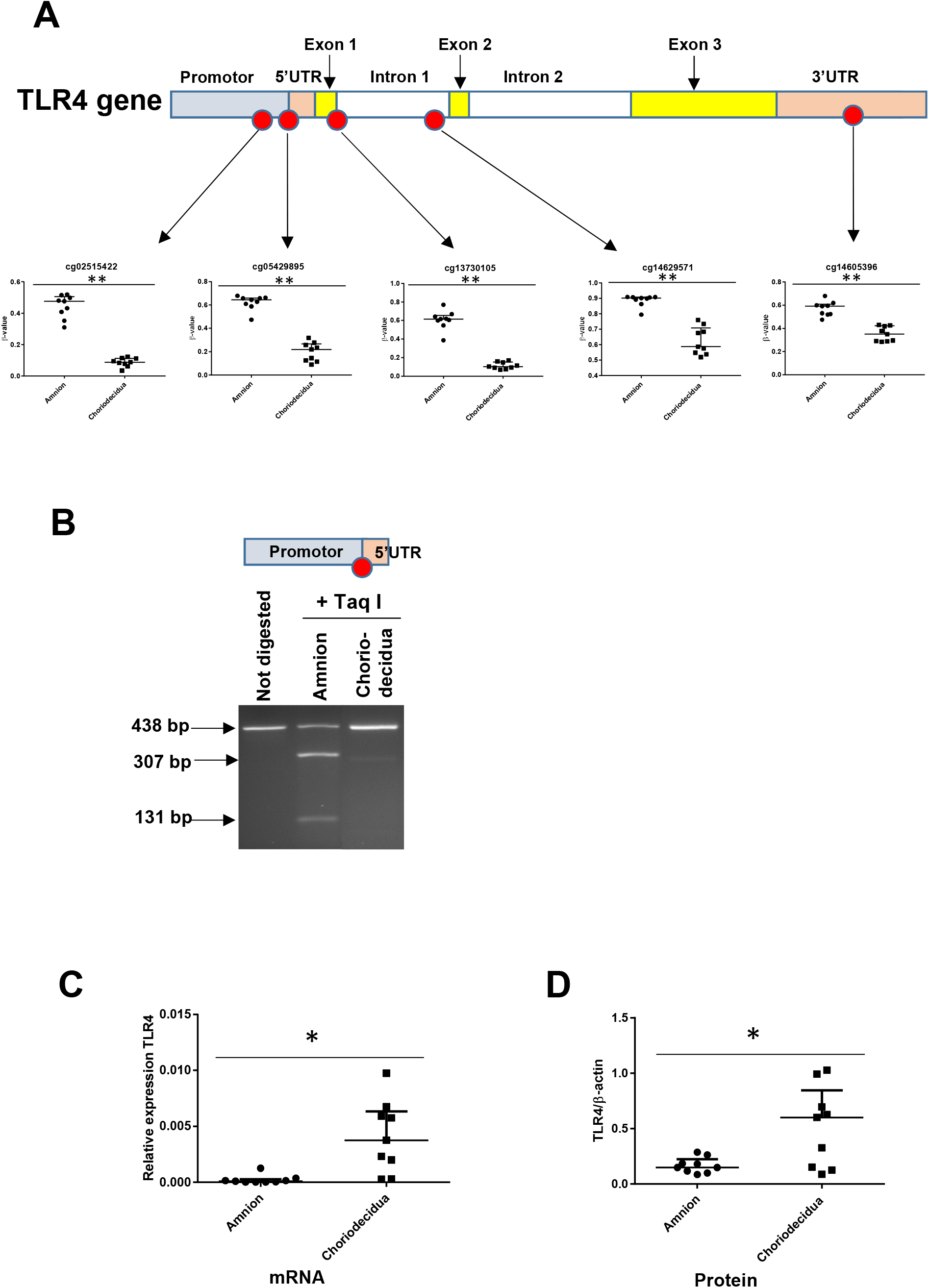
The expression level of the TLR4 gene is related to its cytosine methylation level in the amnion and the choriodecidua. A Top: The median ± interquartile ranges of the DNA methylation levels of the five cg probes on the TLR4 gene in the amnion and choriodecidua ZAMs. Each dot represents the individual β-value for one patient (n=9). The probe set with significant differential methylation between the amnion and choriodecidua (Wilcoxon matched-pairs signed rank test) is designated by an asterisk (** p-value < 0.01). Bottom: Control of the difference of methylation for the cg 05429895 between the amnion and the choriodecidua after digestion with Taq I PCR products (438 bp) obtained after DNA bisulfite treatment. This probe, methylated in the amnion, was sensitive to Taq I and gave two fragments after digestion: 307 bp and 131 bp. B Relative expression of TLR4 transcripts for the nine samples of choriodecidua ZAM were significantly higher than for the amnion ZAM (Wilcoxon matched-pairs signed rank test * p-value < 0.05). C The TLR4 protein was significantly overexpressed for the nine samples in the choriodecidua compared to the amnion (Wilcoxon matched-pairs signed rank test * p-value < 0.05).

### The tissue specificity of TLR4 regulation could be due to miRNA action

Because gene expression can be also regulated post-transcriptionally by the action of mi-RNA, it would be interesting to perform an *in-silico* analysis of differentially methylated miRNA (linked to TLR4) between the amnion and the choriodecidua. The results showed that two of them (let-7a-2 and miR-125b-1) could target the 3’UTR region of TLR4 mRNA (Figure 5A). Focusing on these two, all the cytosines studied on the methylomic chip were situated in the 5’ upstream sequence of miRNA. Five of six were statistically significant for over-methylation in the choriodecidua compared to the amnion for miR-125b-1 and for the unique one in, let-7a-2 (Figure 5B up and bottom panel). Using the same samples that were previously used for the quantification of mRNA TLR4, the expression of each pri-miRNA by the qRT-PCR was checked. As expected, an overexpression of pri-miR-125b-1 was observed in the amnion compared to the choriodecidua. A weak basal expression level below the detection limit of the qPCR assay for pri-let-7a-2 did not allow for obtaining results that could reasonably be analyzed (Figure 5C).

**Figure 5:**
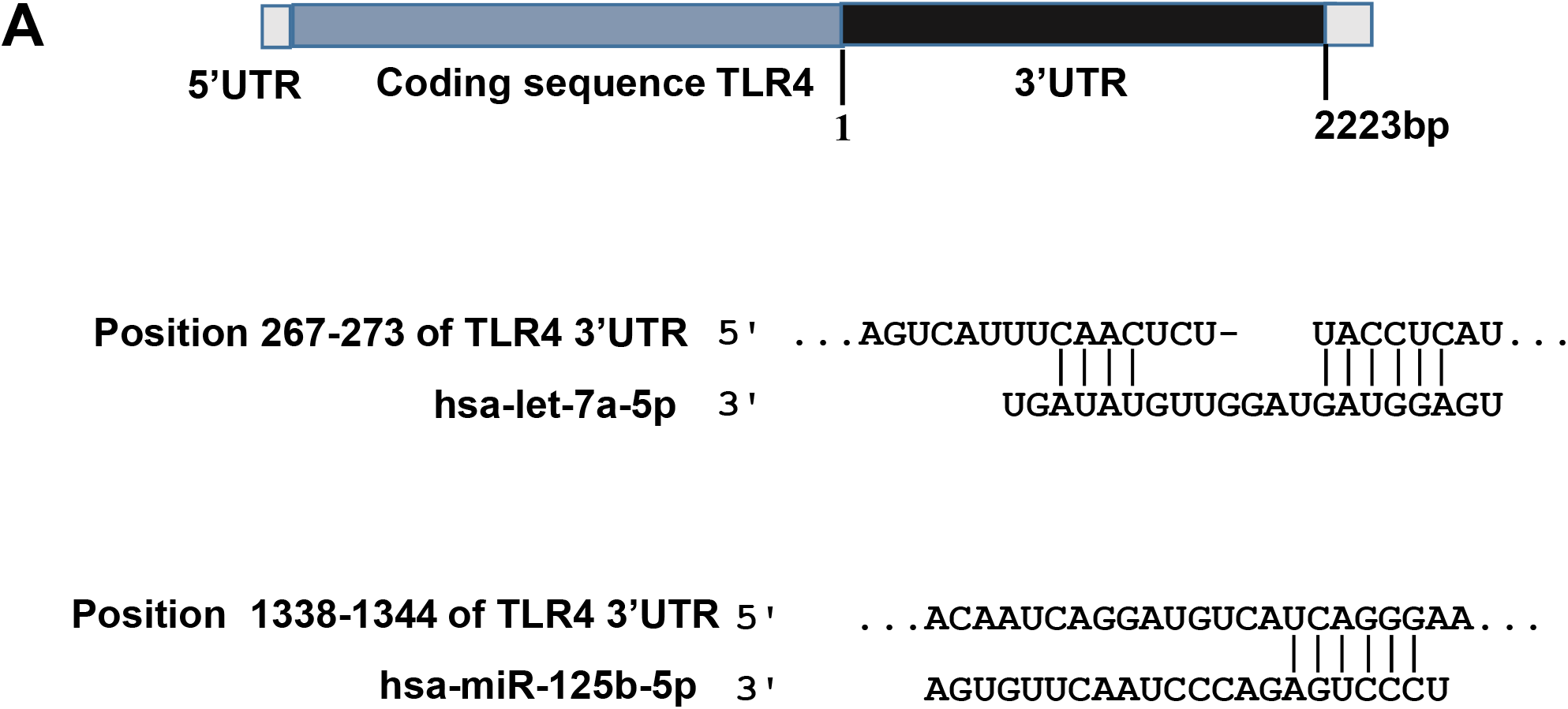
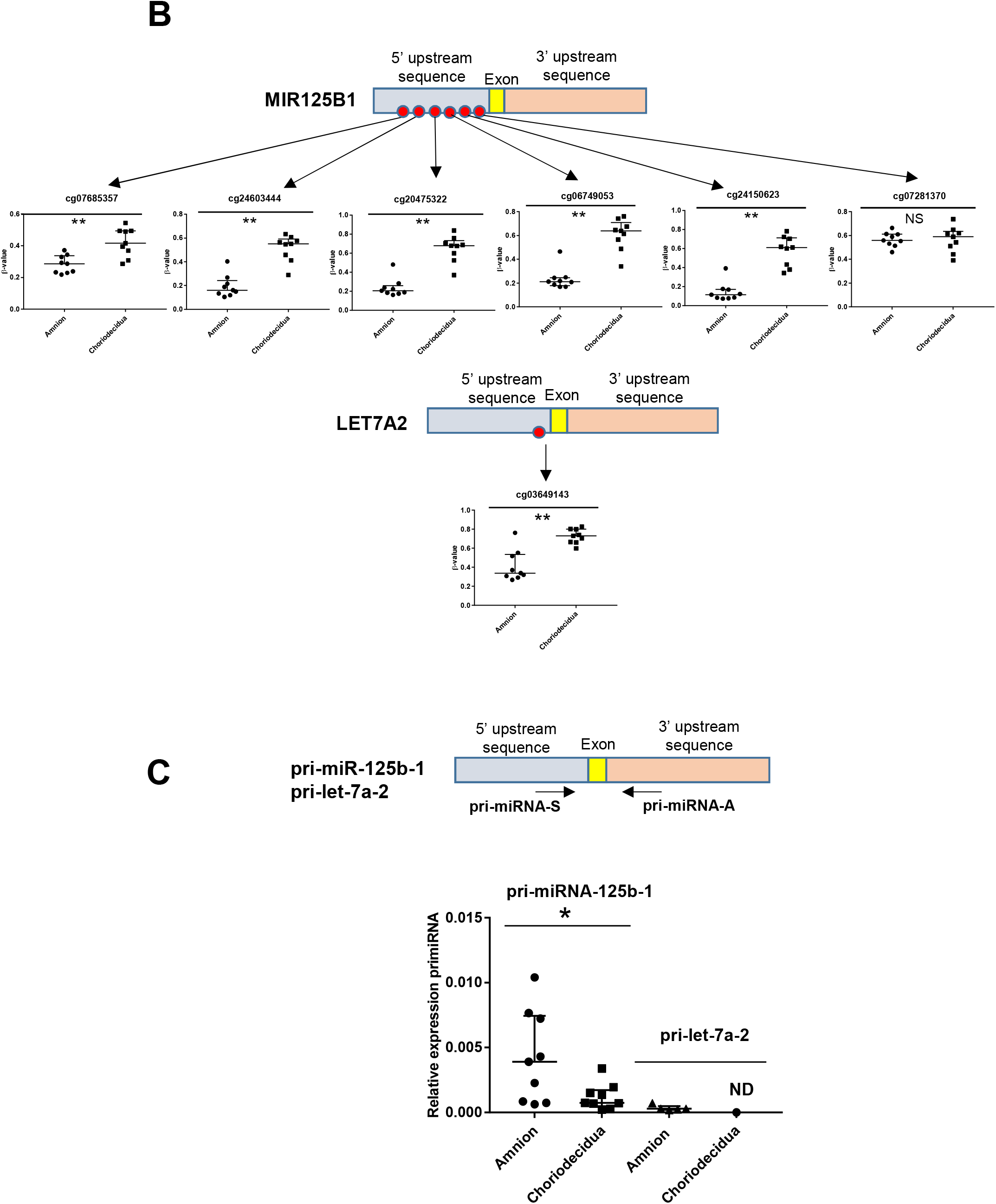
Two miRNAs potentially targeting the human 3’UTR-TLR4 are differentially methylated in the ZAM between the amnion and the choriodecidua. A *In-silico* computational target prediction analysis using TargetScan of the human 3’UTR-TLR4. This zone may be targeted by gene coding for MIR125B1 and LET7A2. B The median ± interquartile ranges of the DNA methylation cg probe levels for the MIR125B1 (top) and LET7A2 (bottom) genes for the nine samples in the amnion and choriodecidua ZAM. Each dot represents the individual β-value for one patient. The probe set with significant differential methylation between the amnion and choriodecidua (Wilcoxon matched-pairs signed rank test) is designated by an asterisk (** p-value < 0.01, NS = not significant). C The relative expression of pri-miR-125b-1 (n=9) and pri-let-7a-2 (n=5) was determined. qRT-PCR experiments performed for each zone, and tissue demonstrated that pri-miR-125b-1 was significantly overexpressed in the amnion compared to the choriodecidua ZAM, as expected from the differential methylation status (Wilcoxon matched-pairs signed rank test * p-value < 0.05). The pri-let-7a-2 amount could not be rigorously determined (ND) in the choriodecidua.

### let-7a-2 and miR-125b-1 target the 3’-UTR-TLR4

We first established that in the human cell line HEK293 (classically used in a gene reporter assay for testing miRNA targets), both miRNAs could decrease the amount of luciferase by targeting the 3’-UTR region of TLR4 mRNA (Figure 6A). To choose a better cellular model to test this miRNA action linked with the fetal membrane environment, the relative amount of TLR4 mRNA was determined in various human amniotic cells. Figure 6B demonstrates that AV3 (one of the three tested amniocyte cell lines, compared to FL and WISH) was the most efficient, and it was therefore used for the following experiments. For *in vitro* or physiological models, AV3 and primary amniotic epithelial cells were respectively used to confirm that both miRNAs clearly targeted TLR4 mRNA or protein accumulation (Figure 6C and 6D). For the AV3 cell line, a significant decrease of the mRNA amount was observed from 12 h after the transfection of let-7a-2 and miR-125b-1 or the co-transfection of let-7a-2+ miR-125b-1. This effect persisted only for let-7a-2 at 24 h, confirming the quick action of this cellular system. Furthermore, after 24 h of transfection, a decrease in the TLR4 protein amount was significantly demonstrated for let-7a-2 and after 48 h for miR-125b-1 or both transfections of let-7a-2 + miR-125b-1. The use of primary cells confirmed this physiological action with a decrease in the TLR4 mRNA amount at 48 h after the transfection of miR-125b-1 and co-transfection of let-7a-2 + miR-125b-1 and after 72 h for the TLR4 protein for the three conditions.

**Figure 6:**
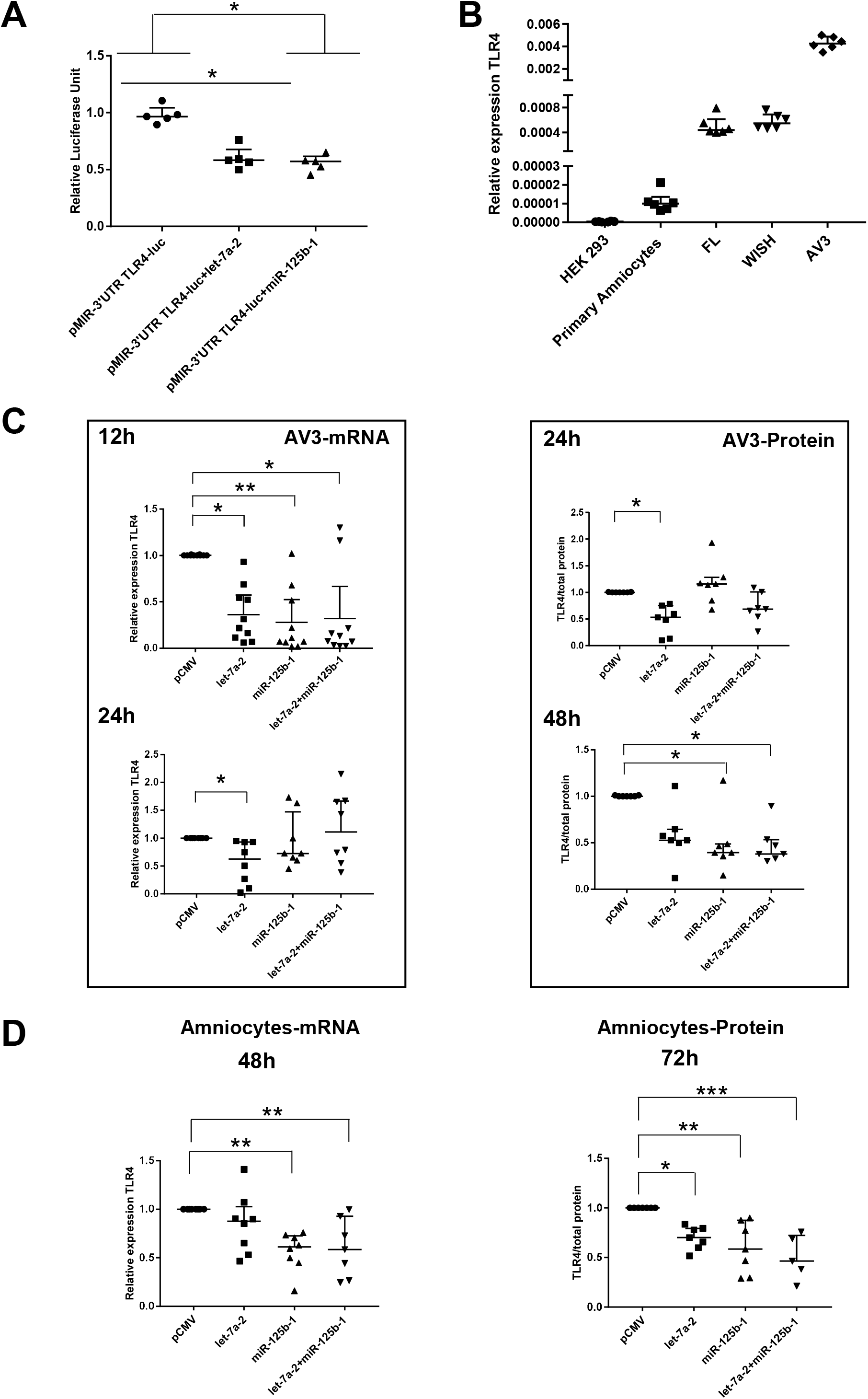
miR-125b-1 and let-7a-2 target the human 3’UTR-TLR4 and decrease TLR4 expression. A Targeting of miR-125b-1 and let-7a-2 to the human 3’UTR of TLR4 mRNA using a Luciferase Reporter Gene Assay depending on human 3’UTR-TLR4 (pMIR-3’UTR TLR4-luc). HEK293 cells co-transfected for 48 h with this construction and expressing plasmid of human pCMV, pre-mir125b1, or let7A2 (n=5). Luciferase activity was normalized with the pRL-TK-Renilla-luciferase level (median ± interquartile range, Kruskal-Wallis test with Dunn’s multiple comparison test, * p-value < 0.05). The results showed that both miRNAs were able to target the 3’-UTR zone of the TLR4 gene and to decrease luciferase quantity. B Determination of endogenous TLR4 expression levels in human cell lines (HEK293 from embryonic kidney, and FL, WISH, and AV3 from Amniocytes) and in primary Amniocyte cells quantified by qRT-PCR (n=6). C Effects of miR-125b-1, let-7a-2, and the combination miR-125b-1 + let-7a-2 on TLR4 mRNA expression (n=10 for 12h and n=8 for 24h, left) and TLR4 protein (n=7, right) in AV3 cells. Cells were transfected with expressing plasmid of human pCMV, or pre-mir125b1, or let7A2 and miR-125b-1 + let-7a-2 for 12 h and 24 h for mRNA, and 24 h and 48 h for protein (median ± interquartile range, Kruskal-Wallis test with Dunn’s multiple comparison test, * p-value < 0.05, ** p-value < 0.01). D Effects of miR-125b-1, let-7a-2, and the combination miR-125b-1 + let-7a-2 on TLR4 mRNA expression (n=7 at least, left) at 48 h and TLR4 protein (n=5 at least, right) at 72 h in primary Amniocyte cells (median ± interquartile ranges, Kruskal-Wallis test with Dunn’s multiple comparison test, * p-value < 0.05, ** p-value < 0.01, *** p-value<0.001).

## DISCUSSION

Worldwide, preterm birth is a serious medical problem, particularly regarding the long-term consequences throughout the entire life of premature infants. PPROM represents one-third of pregnancies that end prematurely and is found to be principally dependent on an early scale and the kinetic activation of inflammation, signaling a cascade in gestational tissues.

Because the inflammatory mechanisms of preterm and term birth are broadly similar, understanding how human fetal membranes are prepared for their physiological rupture is essential. Studies have been partially documented and establish that a weak zone (ZAM) situated in the cervix zone emerged at the end of the nine month of the gestation period (McLaren *et al*, 1999) as a direct consequence of global layer disorganization and weakening. All these phenomena are key determinants of a rupture and could be directly linked to a common denominator: the regulation and modification of gene expression levels (Romero *et al*, 2006b). For the first time, a complementary high-throughput approach was applied by performing both transcriptomic and methylomic analyses on the same samples. Moreover, the strength of this work is to highlight both the global geographical (ZIM/ZAM) and tissue layer (amnion vs. choriodecidua) differences in terms of DNA methylation and gene expression.

Indeed, our methylomic study established a global profile of fetal membranes on the ZIM and ZAM samples. The number of hypermethylated genes in the amnion or the choriodecidua were more important in the ZAM than in the ZIM and allowed us to perform a GO-term classification focused on the ZAM, where cell adhesion, response to a stimulus, tissue development, and reproductive processes are significantly represented. We found some similarities with the transcriptomic analysis, where overexpressed genes in the choriodecidua ZAM were linked to biological adhesion (important in tissue integrity), regulation of cell proliferation, extracellular matrix organization, and responses to internal and external stimuli.

By cross-analyzing both methods for the ZAM, the link of 105 genes overexpressed in the choriodecidua and hypomethylated in the amnion with a MeSH disease terms (e.g., pregnancy complication) was revealed. Especially for GO terms, these highlighted processes could be directly linked to preparation for parturition through a sterile inflammation cascade. We then turned our attention to TLR4, a major mediator of inflammation, for the following reasons. First, this upstream gatekeeper of innate immune activation was the only gene that appeared in each of the considered GO terms, as illustrated in Figure 3A. Second, it is an important partner of many other genes included in our 105 genes list (see Figure 3B). Third, TLR4 was already known to be expressed in the fetal membranes and cervices of animals (Gonzalez *et al*, 2007; Harju *et al*, 2005; Moço *et al*, 2013) and to play a role as a key regulator in a sterile or septic inflammatory reaction in response to aggression by a pathogen-associated molecular pattern (PAMP)/damage-associated molecular pattern (DAMP), leading to PPROM (Chin *et al*, 2016; Li *et al*, 2010; Patni *et al*, 2007; Wang & Hirsch, 2003). Fourth, TLR4 is also known to be involved in parturition at the classical term of pregnancy, activating a sterile inflammation cascade (Choi *et al*, 2012; Wahid *et al*, 2015). Fifth, it could be considered a promising therapeutical target to prevent preterm birth through a better control of its pro-inflammatory signaling.

According to a previous study (Krol *et al*, 2010), and as confirmed by our qPCR and Western blot quantification, TLR4 was more expressed in the choriodecidua than in the amnion (Figure 4B and 4C). This overexpression could be due to a differential methylation and/or transcription factor fixations or could be a consequence of post-transcriptional regulation by miRNAs. miRNAs play crucial regulatory roles in biological and pathological processes. They are well-documented in gestational tissues, such as the placenta, endometrium, and fetal membranes (Doridot *et al*, 2014; Gu *et al*, 2013; Kamity *et al*, 2019; Montenegro *et al*, 2007; Wang *et al*, 2016). They have actually begun to be considered potential biomarkers for pregnancy pathologies (Cretoiu *et al*, 2016), and studies have already shown that some miRNA expressions decrease in chorioamniotic membranes with gestational age uninfluenced by labor (Montenegro *et al*, 2007, 2009). We found two miRNAs (mir-125b-1 and let-7a-2) that were hypermethylated in the choriodecidua compared to the amnion that could potentially target TLR4 mRNA in the amnion to decrease its expression level. They were never known to be able to specifically target TLR4, leading us to demonstrate the consequences of their transfection on the quantification of the TLR4 mRNA and protein on the cell line AV3 or primary amniotic epithelial cells using the luciferase 3’UTR of the TLR4 reporter gene.

Surprisingly, these two miRNAs are part of a miR-100-let-7a-2 cluster host gene, also known as MIR100HG. This cluster has never been studied in relation to the placenta or fetal membrane environment, unlike others, such as C14MC, C19MC, and miR-371-3, whose expressions change during pregnancy between the whole and terminal villi (Gu *et al*, 2013; Morales-Prieto *et al*, 2013). In humans, 10 mature let-7s are synthetized and are implicated in different physiological and pathological events from embryogenesis to adult development, such as inflammatory responses and innate immunity (Roush & Slack, 2008); however, the latter finding could not be linked only to simple TLR4 targeting because both miRNAs were also already known to have implications for innate immunity by influencing not only the expression of interleukins and TNFα but also cell senescence—another well-known phenomenon occurring at the end of fetal membrane life (Iliopoulos *et al*, 2009; Nyholm *et al*, 2014; Schulte *et al*, 2011; Tili *et al*, 2007).

On a global scale, the exhaustive results obtained here provide causal information regarding the implication of inflammation in the physiological rupture of fetal membranes, whereas future studies are needed to exploit all the data accumulated during this work. Nevertheless, by focusing on a unique candidate, TLR4, a well-known actor in the physiological and pathological rupture of membranes, we have outlined the complex molecular process of gene regulation and have proposed a fetal membrane layer-specific model to better understand fetal membrane ruptures (Figure 7). The latter could doubtlessly be extrapolated to other gestational tissues or resident immune cells, such as neutrophils, which are implicated in the amplification of the inflammatory response in late gestation. Moreover, this scheme shows that TLR4 could be an attractive drug target for prolonged gestation and could protect the fetus against inflammatory stress (Robertson *et al*, 2020).

**Figure 7:**
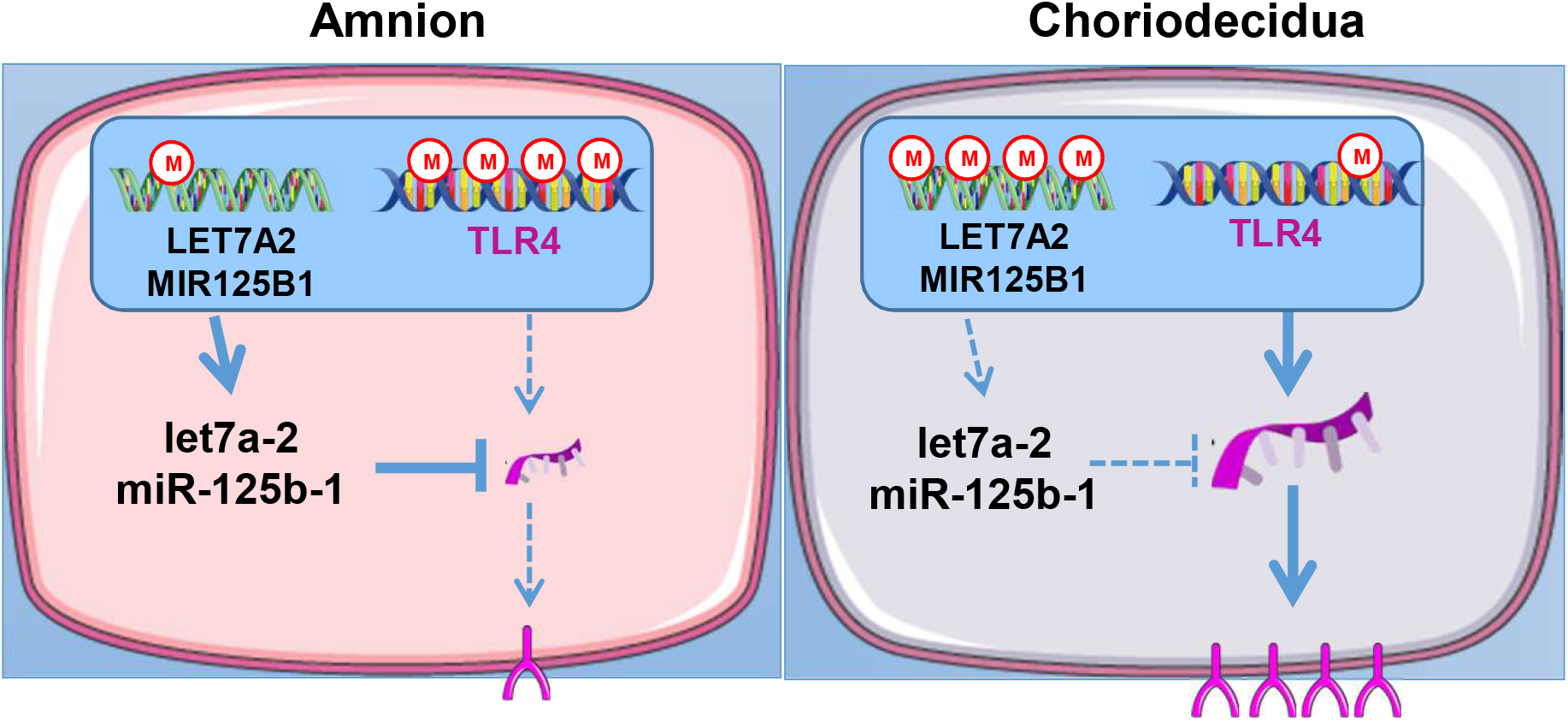
A model for the regulation of TLR4 expression in human fetal membranes is proposed.

All together, these data allow for the expectation of the emergence and promising clinical strategies for pregnant women to prevent PPROM, which will undoubtedly come from results such as ours regarding the physiological preparation of fetal membrane ruptures at term.

## MATERIALS AND METHODS

### Ethics statements

This study was approved by the regional institutional ethics committee, and informed consent was obtained from the participants. The experiments conformed to the principles set out in the World Medical Association Declaration of Helsinki.

### Human fetal membrane collection

Nine fetal membranes were collected at full term before labor and birth by cesarean section (Obstetrics Department, Estaing University Hospital, Clermont-Ferrand, France). All patients were Caucasian and presented no pregnancy pathology (confirmed by macroscopic and microscopic placenta analyses and histological examinations that excluded chorioamnionitis). Supplementary Table S1 describes the patients’ characteristics.

Four samples were collected from each patient (ZAM amnion, ZAM choriodecidua, ZIM amnion, and ZIM choriodecidua) as already described by Choltus *et al* (2020).

### Genome-wide DNA methylation

Total genomic DNA from the amnion and choriodecidua was extracted using the QIAamp^®^ DNA Mini Kit (Qiagen, Courtaboeuf, France) following the manufacturer’s instructions. DNA concentration was determined by Qubit^™^ quantitation (Invitrogen, Thermo Fisher Scientific, Illkirch, France). Five hundred ng of extracted DNA was bisulfite-treated using the EZ DNA Methylation^™^ Kit (Proteigene, Saint-Marcel, France), following the manufacturer’s standard protocol, and individually hybridized using HumanMethylation450 Analysis BeadChip (Illumina, San Diego, California, USA), which allowed for studying the 482,421 cytosines on the human genome. This array was conducted using Helixio (Saint-Beauzire, France). Specific CpG probe methylation differences between the tissues taken from the ZIM and ZAM regions or between the amnion and choriodecidua were analyzed. To study each cytosine, methylation differences were defined as a change in β values (Δβ) between two conditions with p-values less than 0.05 (after applying a Student’s paired t-test), and the Monte Carlo method (Metropolis and Ulam, 1949) was then used to randomly stimulate the minimal (m) and maximal (M) Δβ-value for each chromosome. Such limits permitted us to keep only the cytosine in which Δβ was inferior to M or superior to M. Gene probes meeting the cut-off criteria for each comparison were submitted for Database for Gene Ontology (GO) enrichment (http://go.princeton.edu) in association with the REVIGO web server (http://revigo.irb.hr; as previously described (Supek *et al*, 2011) to identify the biological processes associated with the genes that showed changes in methylation status.

### Human 8x60K expression arrays

Total RNA was extracted using an RNeasy^®^ Mini Kit (Qiagen) following the manufacturer’s instructions. RNA samples were quantified with a NanoDrop^™^ spectrophotometer (ThermoScientific, Thermo Fisher Scientific, Illkirch, France). The RNA integrity was evaluated using a 2100 Bioanalyzer (Agilent Technologies, Les Ullis, France) and an RNA 6000 Nano Assay Kit. The mean of the RNA integrity number (RIN) for all samples was 8.41.

A microarray (Sure Print G3 Human GE 8x60K) was performed by Helixio (Saint-Beauzire, France) in accordance with Agilent Technologies. The data were analyzed with Genespring GX 12.0 (Agilent Technologies).

The resulting gene lists from each pair-wise comparison (ZIM vs. ZAM and amnion vs. choriodecidua) were respectively filtered for the genes that showed 2.8-fold changes at p < 0.01 using the Student–Newman–Keuls test with Benjamini–Hochberg corrections for false discovery rates. Raw data were analyzed with the R (http://R-project.org) and Bioconductor (http://www.bioconductor.org) software. A generic GO term finder (http://go.princeton.edu) and REVIGO (http://revigo.irb.hr/) with up- and downregulated probes were used to analyze the biological process of gene ontology (GO) enrichment, and GePS Genomatix software (Release 2.4.0 Genomatix^®^ Software GmbH, Munich, Germany) was used for the analysis of MeSH diseases (p < 0.01).

### TLR4 immunofluorescent staining

Immunohistochemistry experiments performed on ZAM cryosections taken from the same samples (8 μm) were fixed in paraformaldehyde (4 % in PBS); blocked with PBS-1X, Triton 0.1 %, and SVF 10%; and incubated overnight with a monoclonal mouse anti-TLR4 antibody diluted 1:400 in PBS (Sc-293072, Santa-Cruz Biotechnology, Dallas, Texas, USA). A secondary antibody (donkey anti-mouse coupled with Alexia 488 Ig G [H+L]; Life Technologies, Thermo Fisher Scientific) diluted at 1:300 was incubated for 2 h on slides. Following Hoechst nuclear staining, the samples were examined under a LSM510 Zeiss confocal microscope (Zeiss, Oberkochen, Germany). For negative controls, sections were incubated without a primary antibody.

### Stimulation of fetal membrane explants by Lipopolysaccharide (LPS)

Amnion and choriodecidua explants (2 cm^2^) were placed in six-well plates with 2 ml of DMEM/F12 (Gibco, Thermo Fisher Scientific) supplemented with 10 % fetal bovine serum (FBS, GE Healthcare, Vélizy-Villacoublay, France), 100 units/ml of penicillin, 100 µg/ml of Streptomycin, and 0.25 µg/ml of Amphotericin B (Hyclone, Thermo Fisher scientific) for 24 h. The medium was then removed and replaced with serum-free DMEM/F12. The explants were treated (or left untreated) with LPS from E.coli (O55:B5) at 0.5 µg/ml (Sigma-Aldrich Chimie, St. Quentin Fallavier, France) for 24 h. The treated explants and cell-free culture medium were collected and stored at -80 °C.

### Cytokine analysis

Secreted IL-6 and TNFα cytokines were measured in an amnion and choriodecidua-conditioned medium using a Bio-Plex Pro^™^ Human Cytokine 27-plex Assay (Bio-Rad, Marnes-la-Coquette, France) as recommended by the manufacturer along with Luminex technology. The results were standardized with the total protein concentration using a Pierce^™^ BCA Assay (Thermo Fisher Scientific).

### Bisulfite conversion and combined restriction analysis

Total DNA from the amnion and choriodecidua explants was bisulfite-treated using an EZ DNA methylation kit (Zymo Research, Proteigene). The converted DNAs were used as templates for the PCR reaction using specific primers of promoter TLR4 (containing the cytosine cg 05429895; promot-F: 5’ TTTAGAGAGTTATAAGGGTTATTT 3’, and promot-R: 5’ CTAACA TCATCCT CACTACTTC 3’). The PCR was performed using HotStart DNA Polymerase (Qiagen), and the products were digested with Taq I (New England Biolabs, Evry, France) that could recognize CpGs. Digested products were analyzed on polyacrylamide gel.

### Cell cultures

Primary amniocyte cells were collected from the amnion after trypsination and were cultured on six-well plates coated with bovine collagen type I/III (StemCell Technologies, Saint-Egrève, France) as described previously (Marceau *et al*, 2006; Prat *et al*, 2015). Human embryonic kidney (HEK) 293 cells were grown in Dulbecco’s Modified Eagle Medium (DMEM; Gibco) supplemented with 10 % fetal bovine serum (FBS, GE healthcare), 4mM glutamine (Gibco), 1mM sodium pyruvate (Hyclone), 1x non-essential amino acids (Gibco), 100 units/ml of penicillin, 100 µg/ml of Streptomycin, and 0.25 µg/ml of Amphotericin B (Hyclone).

Human epithelial amnion cells (AV3 cells, ATCC-CCL21; FL, ATCC-CCL62; WISH, ATCC-CCL25) were cultured in DMEM/F12 (Gibco) supplemented with 10 % fetal bovine serum (FBS, GE healthcare), 4 mM glutamine (Gibco), 100 units/ml of penicillin, 100 µg/ml of Streptomycin, and 0.25 µg/ml of Amphotericin B (Hyclone).

### Quantitative RT-PCR

Total RNA was isolated from the amnion, choriodecidua, human epithelial amnion cells (AV3, FL, and WISH), and primary amniocyte cells using an RNeasy^®^ Mini Kit (Qiagen) with DNAse I digestion as described in the manufacturer’s protocol. After quantification with a NanoDrop^™^ spectrophotometer (Thermo Fisher Scientific), cDNA was synthesized from 1μg of RNA using a SuperScript^™^ III First-Strand Synthesis System for RT-PCR (Invitrogen, Thermo Fisher Scientific). Quantitative RT-PCR reactions were performed with a LightCycler^®^ 480 (Roche Diagnostics, Meylan, France) using Power SYBR^®^ Green Master Mix (Roche) and specific primers (described in supplementary Table S2). Transcripts were quantified using a standard curve method. The ratio of interest (transcript/geometric) mean of two housekeeping genes (RPLP0 and RPS17) was determined. The results were obtained from at least six independent experiments, and all the steps followed the MIQE guidelines (Bustin *et al*, 2009).

### Western blot

Primary amniocyte and AV3 cells were lysed in a RIPA buffer (20 mM Tris-HCl pH = 7.5, 150 mM NaCl, 1 % Nonidet P40, 0.5 % sodium deoxycholate, 1mM EDTA, 0.1 % SDS, 1x Complete Protease inhibitor cocktail (Roche) for 30 min at 4 °C. The amnion and choriodecidua samples were homogenizated as described (Choltus *et al*, 2020). The protein concentration of the supernatant was determined using a Pierce™ BCA Assay Kit (Pierce). Forty µg of denaturated proteins were subjected to a Western blot analysis after 4–15 % MiniPROTEAN^®^ TGX StainFree^™^ gel electrophoresis (Bio-Rad) followed by probing antibodies against TLR4 or β-Actin (TLR4:1:400, Sc-293072, Santa-Cruz Biotechnology, β-Actin:1:10,000, MA1-91399, Thermo Fisher Scientific). The signal was detected with a peroxidase-labeled anti-mouse antibody at 1:10,000 (Sc-2005, Santa Cruz Biotechnology) and visualized with ECL or ECL2 Western blotting substrate (Pierce) using a ChemiDoc^™^ MP Imaging System and Image Lab™ software (Bio-Rad). The results were obtained from at least seven independent experiments and the relative TLR4 ratio was expressed in function of β-Actin or total protein loaded by well.

### 3’UTR-hTLR4 pMIR REPORT^™^ luciferase plasmid

The 3’UTR hTLR4 (2,223 bp, NM 138554) was amplified from 100 ng of genomic amnion DNA with 3’UTR-TLR4 primers containing Spe I restriction sites to facilitate subcloning: forward GAGA*ACTAGT*AGAGGAAAAATAAAAACCTCCTG and reverse GAGA *ACTAGT*TTGATATTATAAAACTGCATATATTTA. The PCR was performed with Phusion^®^ high-fidelity DNA polymerase (New England Biolabs) according to the manufacturer’s instructions. After purification and digestion with Spe I (New England Biolabs), the fragment was subcloned into the Spe I site of the pMIR-REPORT^™^ luciferase vector (Ambion, Thermo Fisher scientific) to generate the construct pMIR-3’UTR-hTLR4. The insert sequence was checked by sequencing (Eurofins Genomics, Ebersberg, Germany).

### *In-silico* miRNA analysis

Putative miRNAs targeting the 3’UTR-TLR4 target gene miRNA were screened using the public database miRWalk with TargetScan, RNA22, and miRanda.

### Dual-Luciferase^®^ Reporter assays

HEK293 cells were cultured in six-well plates, transiently transfected at an 80–90 % confluence using a Lipofectamine^®^ 3000 Transfection Kit (Invitrogen, Thermo Fisher Scientific). Each well received 100 ng of the pMIR-REPORT^™^ Luciferase vector containing the 3’UTR-hTLR4 (without a negative control) in combination with 1µg pCMV-MIR or pCMV-pre-mir125b1 (OriGene Technologies, Rockville, Maryland, USA) or let7A2 generated in the laboratory and with 50 ng of pRL-TK Renilla luciferase plasmid (Promega).. Briefly, the pre-let7A2 was amplified from 100 ng of genomic amnion DNA with pre-let7A2 primers containing SgfI / MluI restriction sites to facilitate subcloning: forward GAGA*GCGATCGCTC*GTCAACAGATATCAGAAGGC and reverse GAGA*ACGCGT*AATGCTGCATTTTTTGTGACAATTT. The PCR was performed with Advantage-HD DNA polymerase (Takara Bio, Saint-Germain-en-Laye, France) according to the manufacturer’s instructions. After purification and digestion with SgfI / MluI (New England Biolabs), the fragment was subcloned into the SgfI / MluI site of the pCMV-MIR vector to generate the construct pCMV-pre-let7A2. The insert sequence was checked by sequencing (Eurofins Genomics, Ebersberg, Germany). Luciferase activity was measured using the Dual-Luciferase^®^ Reporter Assay System (Promega, Charbonnières-les-Bains, France) 48 h after transfection according to the manufacturer’s instructions using a Sirius luminometer (Berthold, thoiry, France). Transfection efficiencies were normalized to the Renilla luciferase activities in the corresponding wells and reported to the control condition. Data were extracted from at least three experiments, each performed in triplicate.

### Transfections of primary amniocytes and AV3 cells

Primary amniocytes and AV3 were cultured in six-well plates. At 80–90 % of confluence, the cells were transiently transfected using a Lipofectamine^®^ 3000 Transfection Kit (Invitrogen, thermos Fisher Scientific) according to the protocol with 1µg expressing plasmid of human pCMV pre-mir125b1, pre-let7A2, or a control (pCMV-MIR) purchased from OriGene. After 48 h or 72 h of culturing, total RNA and proteins were respectively collected and used for the qRT-PCR or Western blot experiments. Data were extracted from five experiments.

### Statistical analysis

The results were analyzed using PRISM software (GraphPad Software Inc., San Diego, California, USA). The quantitative data were presented as the medians ± interquartile ranges according to a Shapiro–Wilks test. Non-normally distributed data between the two groups were studied with a Wilcoxon matched-pairs signed rank test for paired samples. To study several independent groups, a Kruskal–Wallis test was performed followed by a Dunn’s multiple comparison test. Values were considered significantly different at p < 0.05 (*), p < 0.01 (**), or p < 0.001 (***) throughout.

## ACKNOWLEDGEMENTS

LB was supported in carrying out this work by the Auvergne-Rhone-Alpes region’s Jeune Chercheur framework. This research was conducted with the scientific support and expertise of Helixio (Saint-Beauzire, France).

## AUTHOR CONTRIBUTIONS

CB, FPC, GC, MR, and CG conducted the experiments and acquired the data.

CB and BP conducted the statistical analyses.

DG, VS, and LB designed the research studies.

CB, VS, and LB wrote the manuscript.

## DATA AVAILABILTY SECTION

-Term W/O labor Fetal membrane methylation status results were submitted into Arrayexpress (http://www.ebi.ac.uk/arrayexpress/experiments/E-MTAB-10520).

-Term W/O labor Fetal membrane mRNA expression results were submitted into Arrayexpress (http://www.ebi.ac.uk/arrayexpress/experiments/E-MTAB-10516)

## TRANSPARENT REPORTING FORM

### FIGURES SOURCE DATA

**Figure 3-source data 1:** Raw data for TLR4 immunofluorescence in the fetal membranes by confocal microscopy.

**Figure 3-source data 2:** Data on IL-6 and TNF-α concentrations in amnion and choriodecidua.

**Figure 4-source data 1:** Data on β-value for cg probes on the TLR4 gene for Figure 4A.

**Figure 4-source data 2:** Uncropped polyacrylamide gel for Figure 4B.

**Figure 4-source data 3:** qRT-PCR data and Western-blots data for TLR4 expression (Figure 4C).

**Figure 5-source data 1:** Data on β-value for cg probes on the MIR125B1 and LET7A2 genes for Figure 5B.

**Figure 5-source data 2:** qRT-PCR data for pri-miR-125b-1 and pri-let-7a-2 (Figure 5C).

**Figure 6-source data 1:** Data on luciferase activity in HEK293 cells transfected with pMIR-3’UTR TLR4-luc and pre-mir125b1 or let7A2 (Figure 6A)

**Figure 6-source data 2:** qRT-PCR data for TLR4 expression for Figure 6B.

**Figure 6-source data 3:** qRT-PCR data and Western-blots data for TLR4 expression in AV3 cells transfected with pre-miRNA (Figure 6C).

**Figure 6-source data 4:** qRT-PCR data and Western-blots data for TLR4 expression in Amniocytes transfected with pre-miRNA (Figure 6D).

## Key resources table

| **Reagent type**<br>**(species) or**<br>**resource** | **Designation** |
| --- | --- |
| Antibody | Monoclonal mouse anti-TLR4 |
| Antibody | Monoclonal mouse anti-β Actin |
| Cell line | Human epithelial amnion cells AV3 |
| Cell line | Human epithelial amnion cells FL |
| Cell line | Human epithelial amnion cells WISH |
| Cell line | Human embryonic kidney (HEK) 293 |
| Commercial<br>assay or kit | QIAamp<sup>®</sup> DNA Mini Kit |
| Commercial<br>assay or kit | EZ DNA Methylation<sup>™</sup> Kit |
| Commercial<br>assay or kit | Bio-Plex Pro<sup>™</sup> Human Cytokine 27-plex Assay |
| Commercial<br>assay or kit | BCA Assay kit |
| Commercial<br>assay or kit | RNeasy<sup>®</sup> Mini Kit |
| Commercial<br>assay or kit | SuperScript<sup>™</sup> III First-Strand Synthesis System |
| Commercial<br>assay or kit | Dual-Luciferase<sup>®</sup> Reporter Assay System |
| Commercial<br>assay or kit | Lipofectamine<sup>®</sup> 3000 Transfection Kit |
| Software,<br>algorithm | GraphPad Prism Software |
| Software,<br>algorithm | GePS Genomatix |
| Software,<br>algorithm | Image Lab<sup>™</sup> software |
| Software,<br>algorithm | miRWalk |
| Software,<br>algorithm | R (CRAN) |
| Software,<br>algorithm | Generic GO term finder |
| Software,<br>algorithm | Bioconductor |
| Software,<br>algorithm | REVIGO |
| Chemical<br>compound, drug | LPS from E.coli (O55:B5) |
| Recombinant<br>DNA reagent | pMIR-REPORT™ luciferase vector |
| Recombinant<br>DNA reagent | pCMV-MIR |
| Recombinant<br>DNA reagent | pCMV-pre-mir125b1 |
| Recombinant<br>DNA reagent | pCMV-pre-let7A2 |
| Recombinant<br>DNA reagent | pRL-TK Renilla luciferase plasmid |

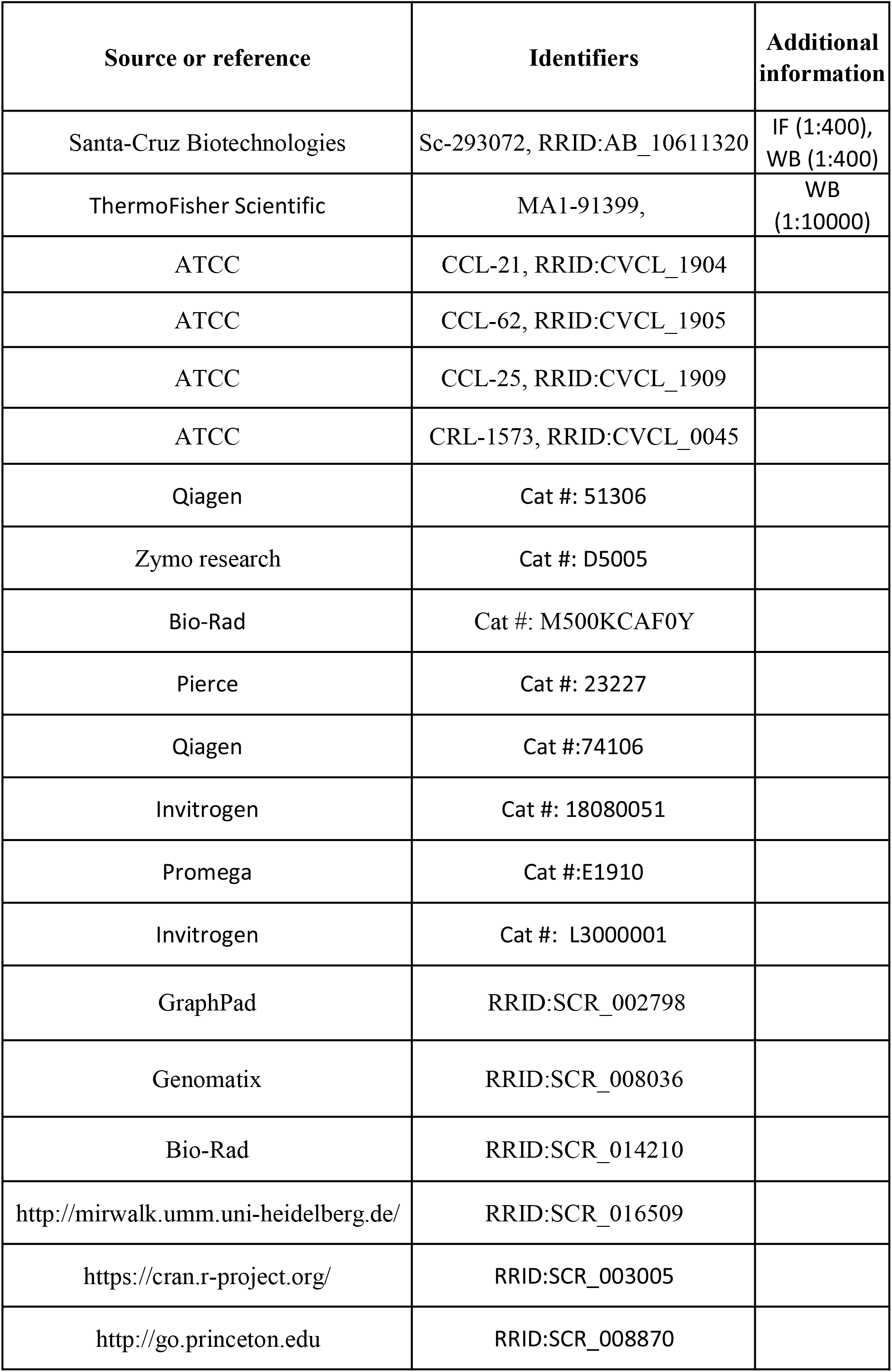
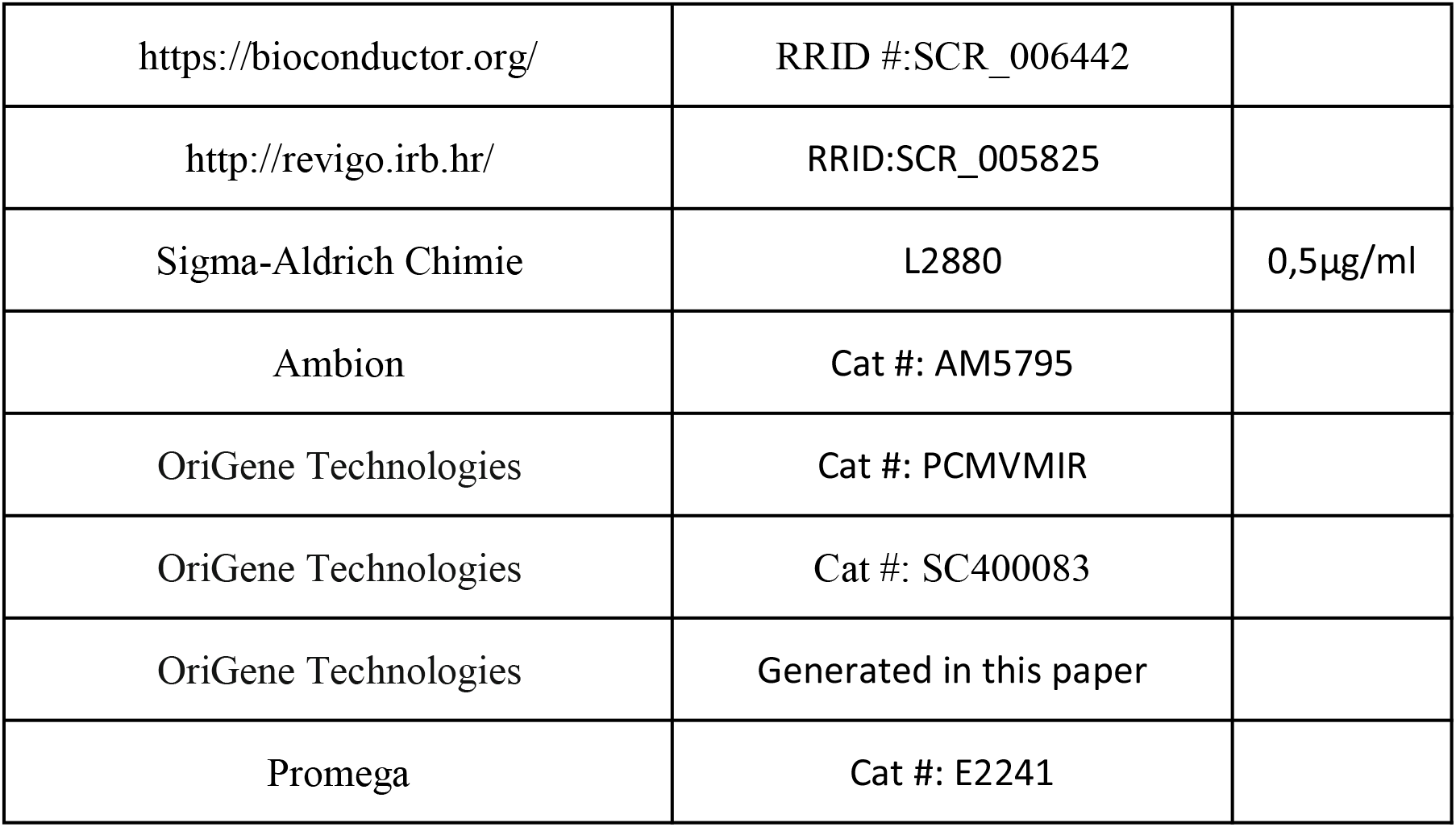

| Source or reference | Identifiers | Additional information |
| --- | --- | --- |
| Santa-Cruz Biotechnologies | Sc-293072, RRID:AB_10611320 | IF (1:400),<br>WB (1:400) |
| ThermoFisher Scientific | MA1-91399, | WB<br>(1:10000) |
| ATCC | CCL-21, RRID:CVCL_1904 |  |
| ATCC | CCL-62, RRID:CVCL_1905 |  |
| ATCC | CCL-25, RRID:CVCL_1909 |  |
| ATCC | CRL-1573, RRID:CVCL_0045 |  |
| Qiagen | Cat #: 51306 |  |
| Zymo research | Cat #: D5005 |  |
| Bio-Rad | Cat #: M500KCAF0Y |  |
| Pierce | Cat #: 23227 |  |
| Qiagen | Cat #:74106 |  |
| Invitrogen | Cat #: 18080051 |  |
| Promega | Cat #:E1910 |  |
| Invitrogen | Cat #: L3000001 |  |
| GraphPad | RRID:SCR_002798 |  |
| Genomatix | RRID:SCR_008036 |  |
| Bio-Rad | RRID:SCR_014210 |  |
| http://mirwalk.umm.uni-heidelberg.de/ | RRID:SCR_016509 |  |
| https://cran.r-project.org/ | RRID:SCR_003005 |  |
| http://go.princeton.edu | RRID:SCR_008870 |  |
| https://bioconductor.org/ | RRID #:SCR_006442 |  |
| http://revigo.irb.hr/ | RRID:SCR_005825 |  |
| Sigma-Aldrich Chimie | L2880 | 0,5µg/ml |
| Ambion | Cat #: AM5795 |  |
| OriGene Technologies | Cat #: PCMV MIR |  |
| OriGene Technologies | Cat #: SC400083 |  |
| OriGene Technologies | Generated in this paper |  |
| Promega | Cat #: E2241 |  |

**Table S1:** Perinatal characteristics of the enrolled patients. F = Female, M = Male. Data are expressed as the mean ± SEM.

| **Perinatal characteristics** | **n=9** |
| --- | --- |
| **Maternal Age (years)** | 32.33 ± 1.76 |
| **Gestational age at delivery (Amenorrhea weeks)** | 39.10 ± 0.08 |
| **Body Mass Index, BMI** | 23.75 ± 0.95 |
| **Child sexe** | 5F, 4M |
| **Birth weight (g)** | 3351 ± 129 |

**Table S2:**
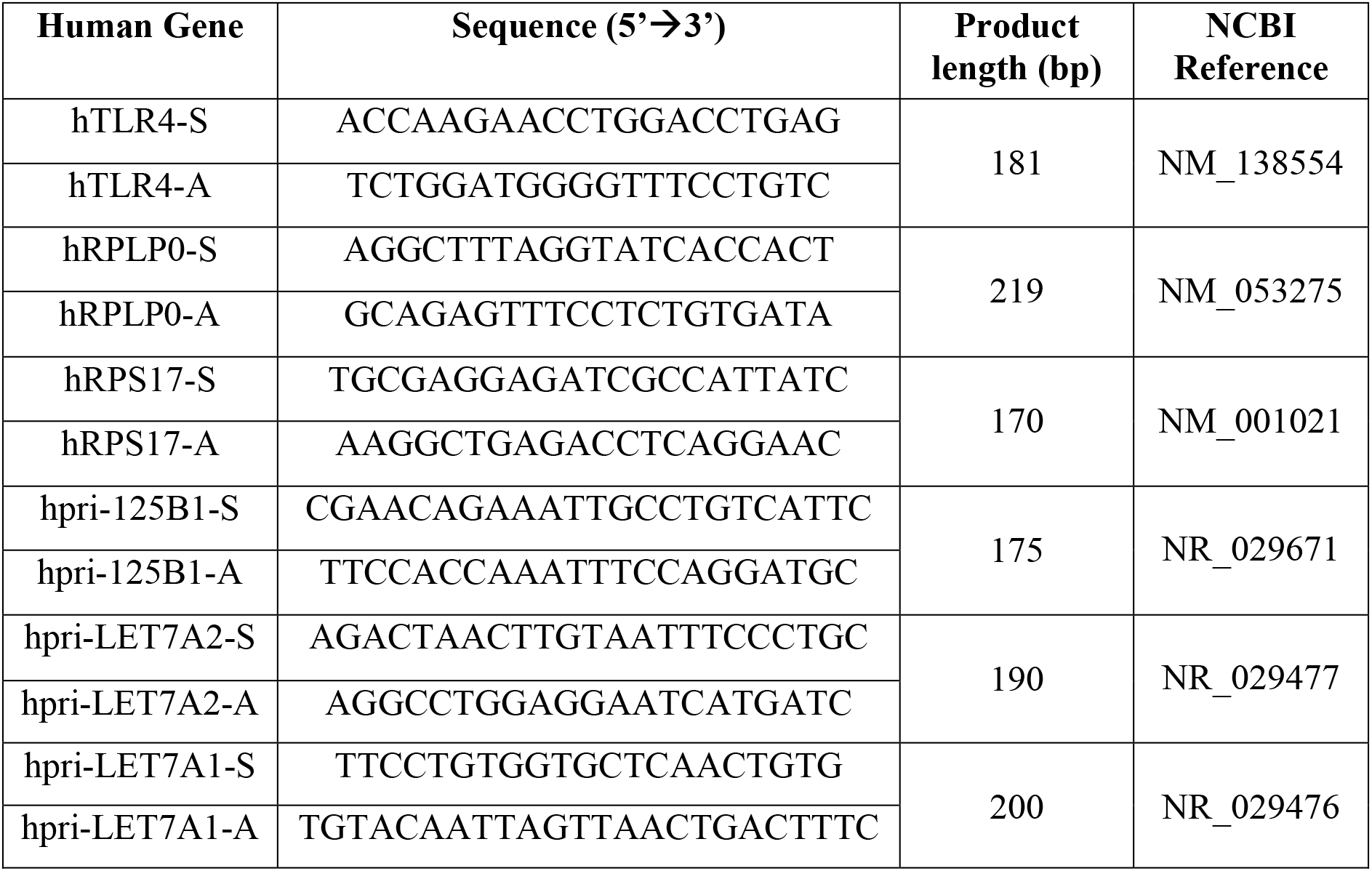
Primer sequences used for the PCR.

**Table S3:**
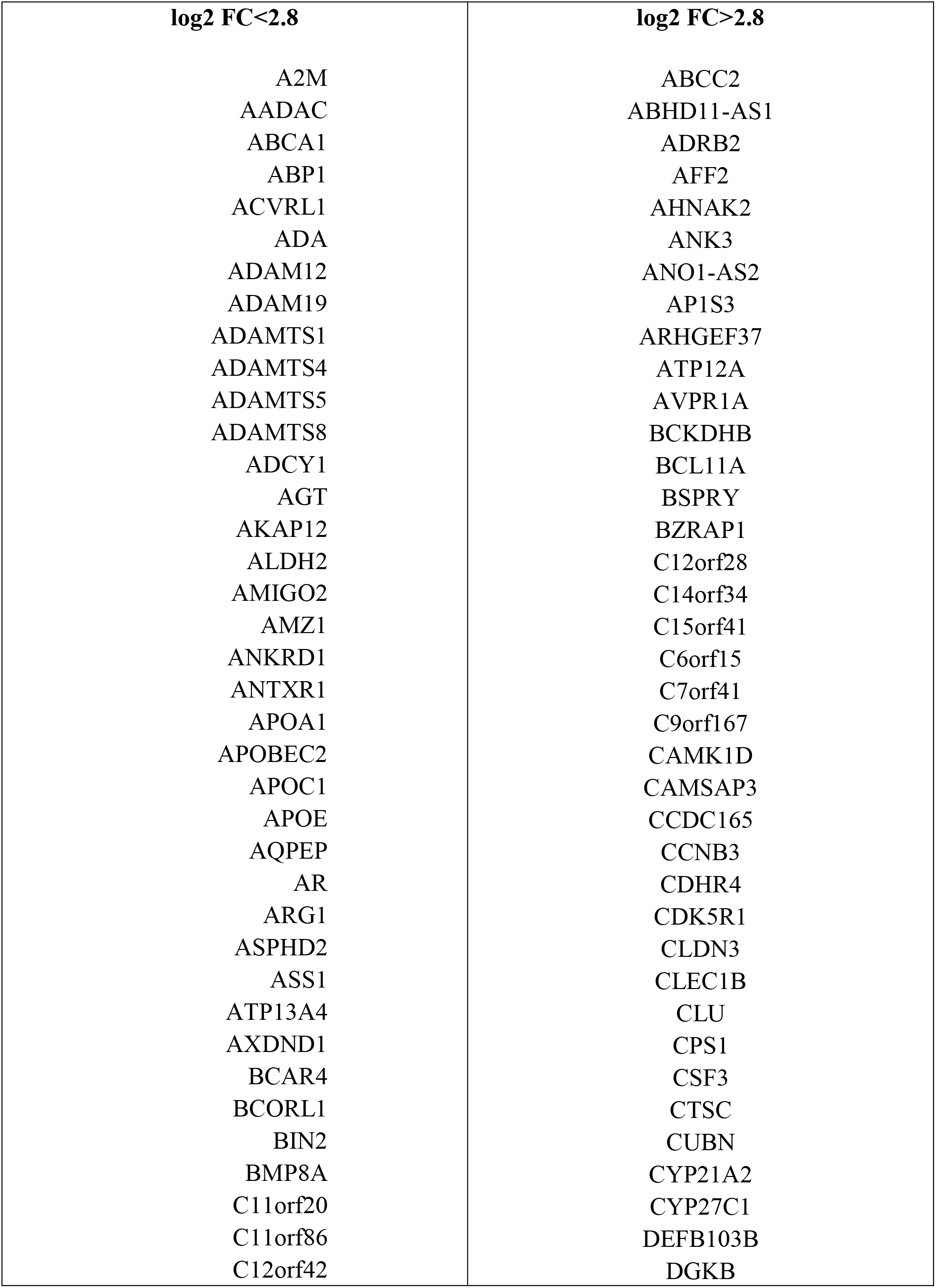
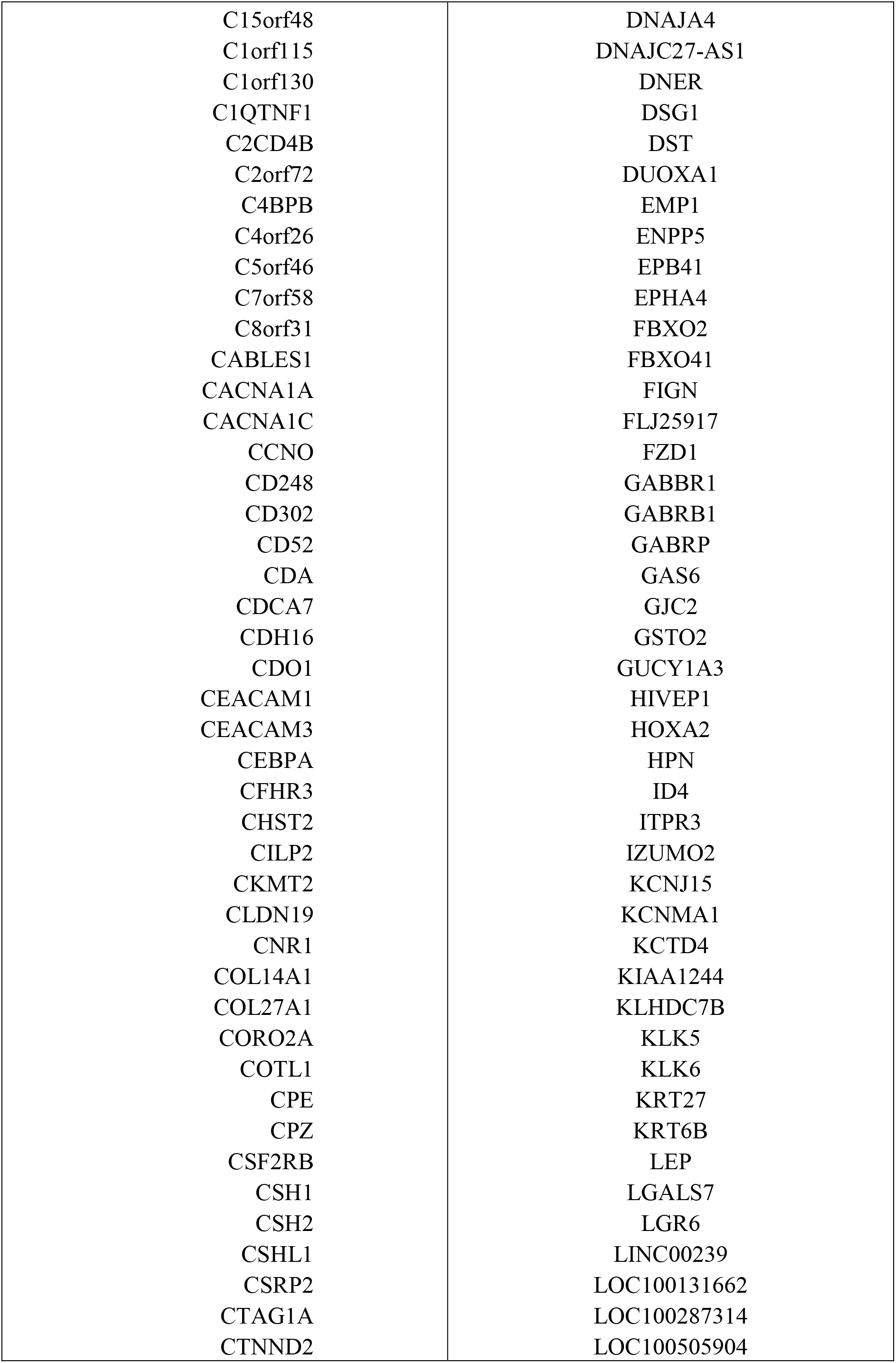
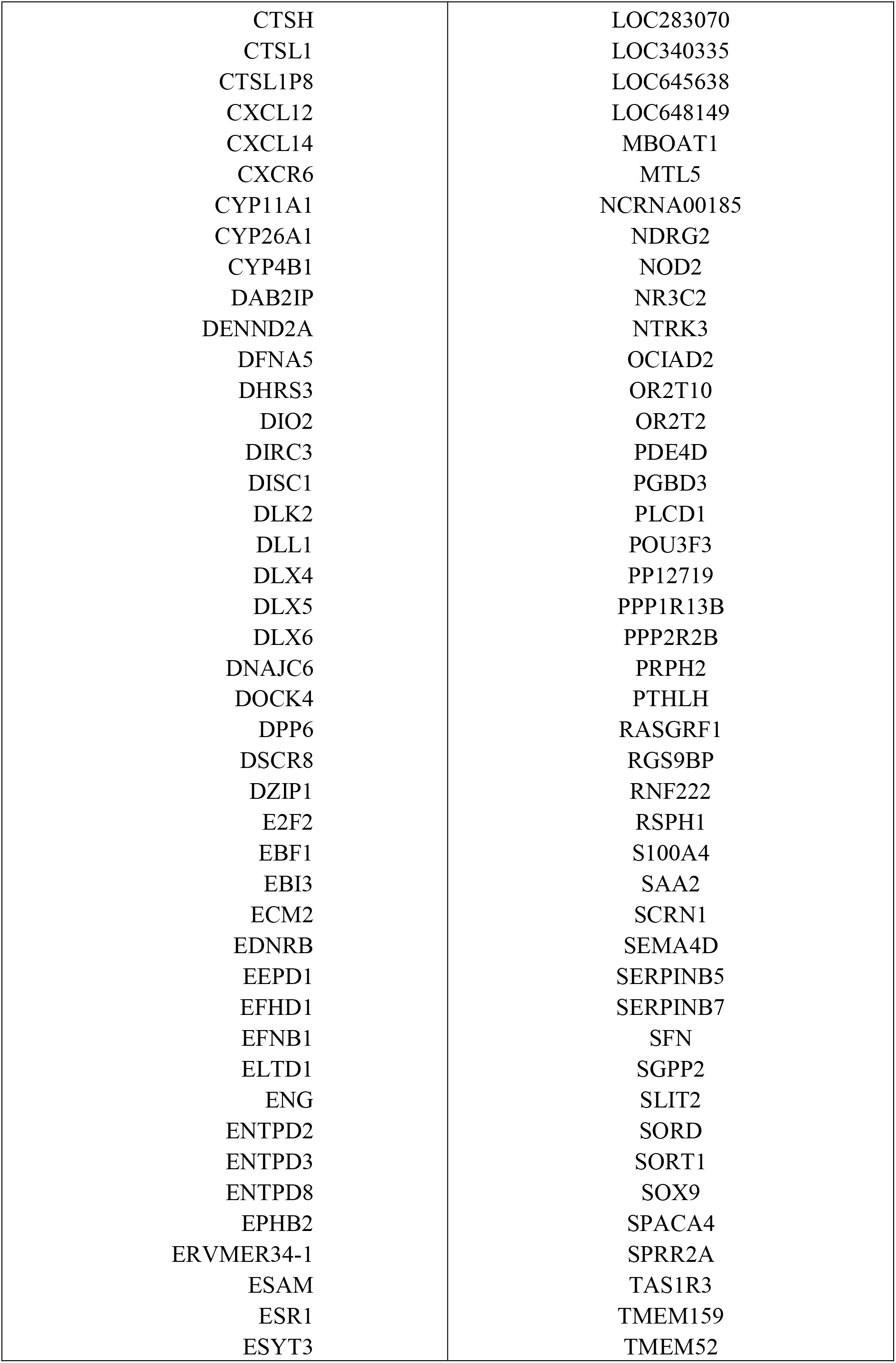
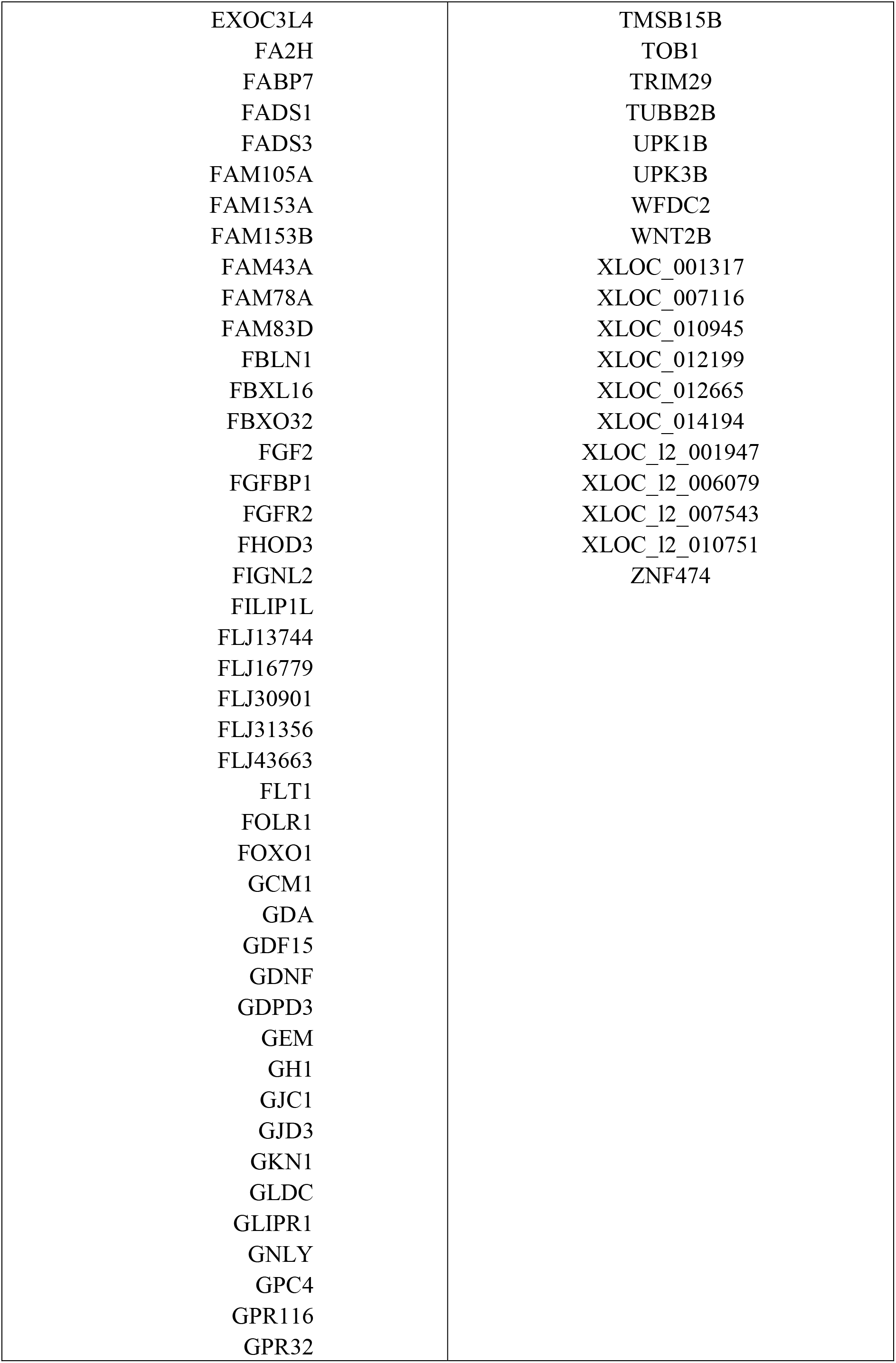
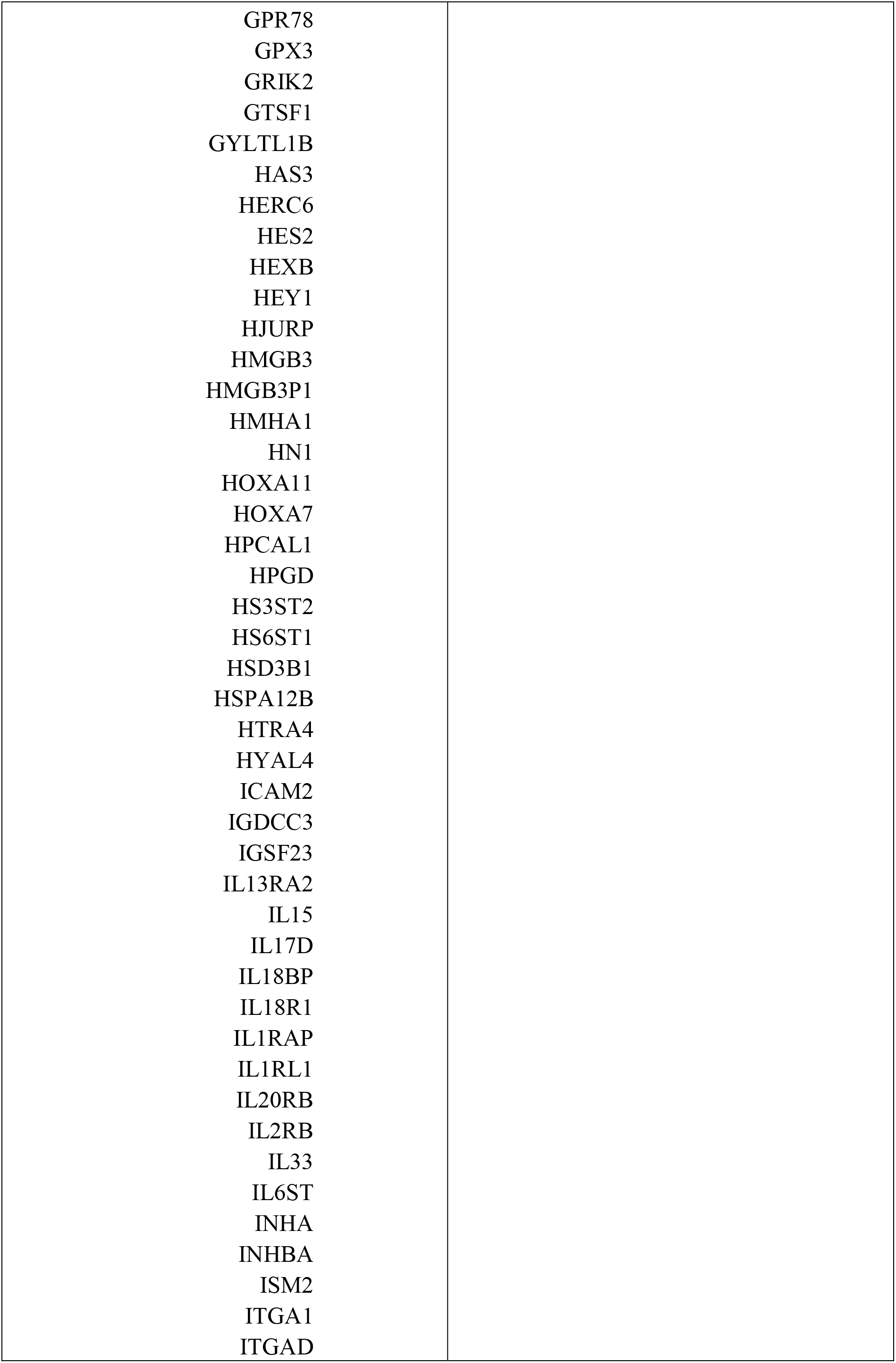
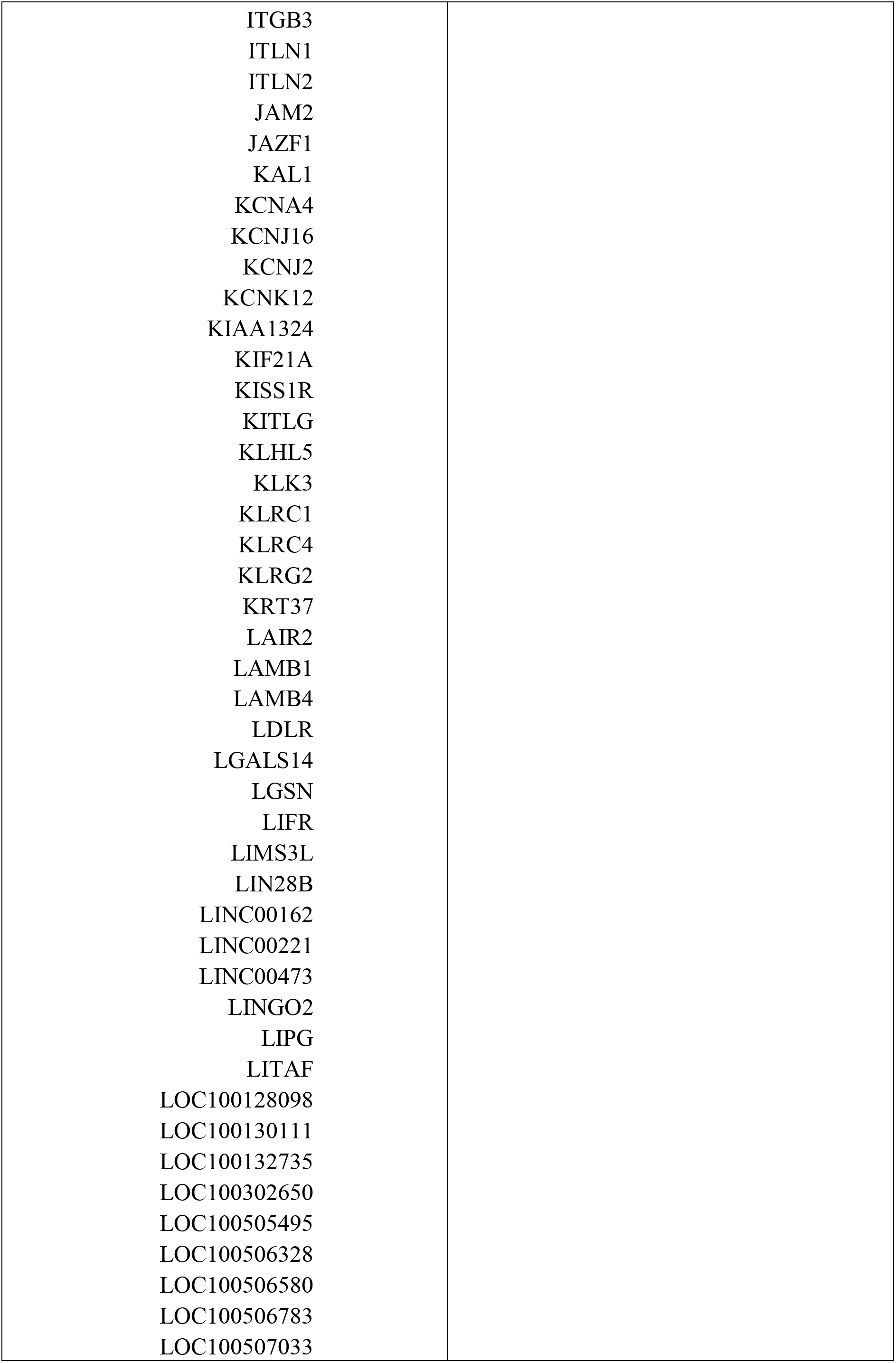
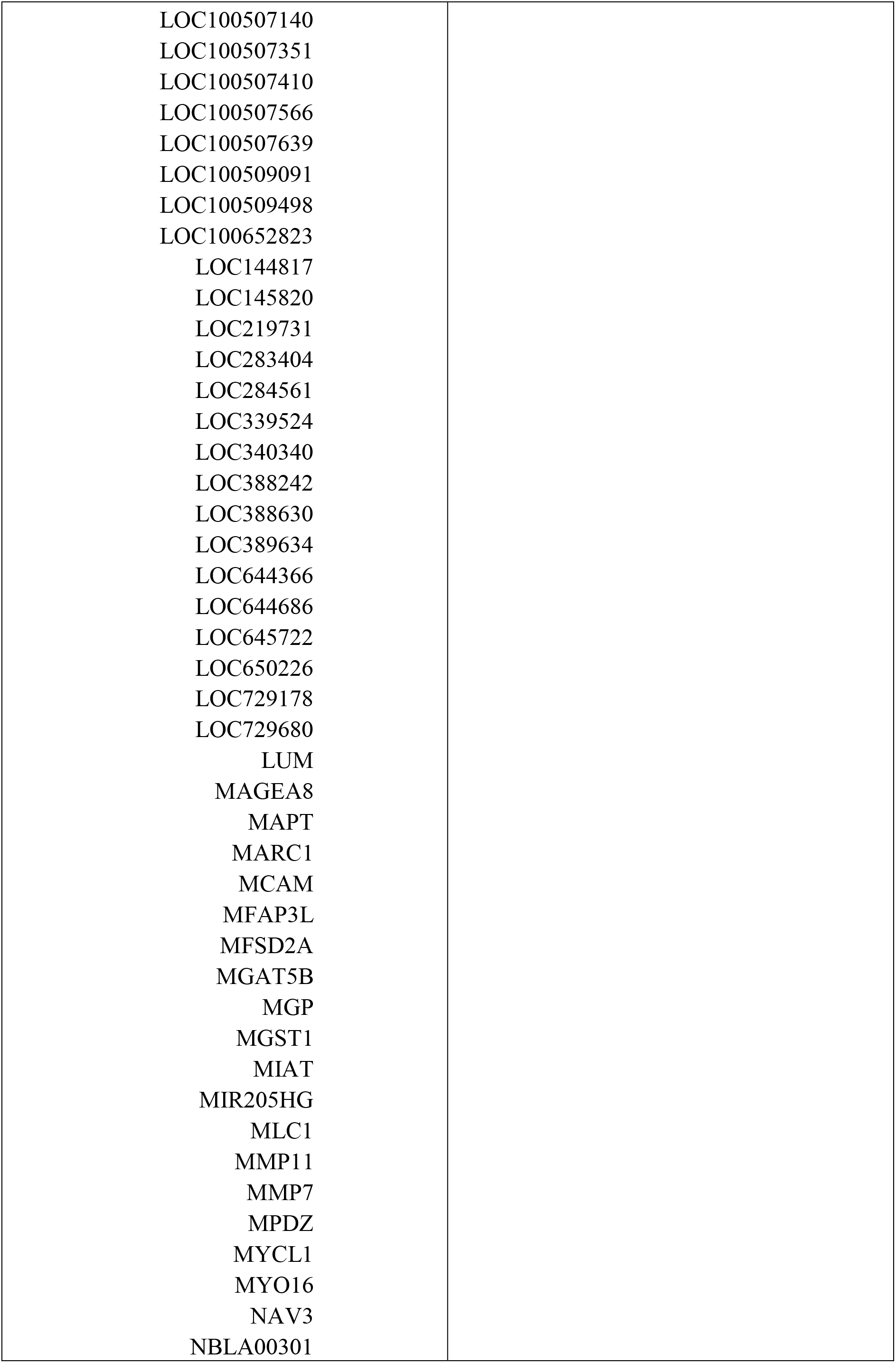
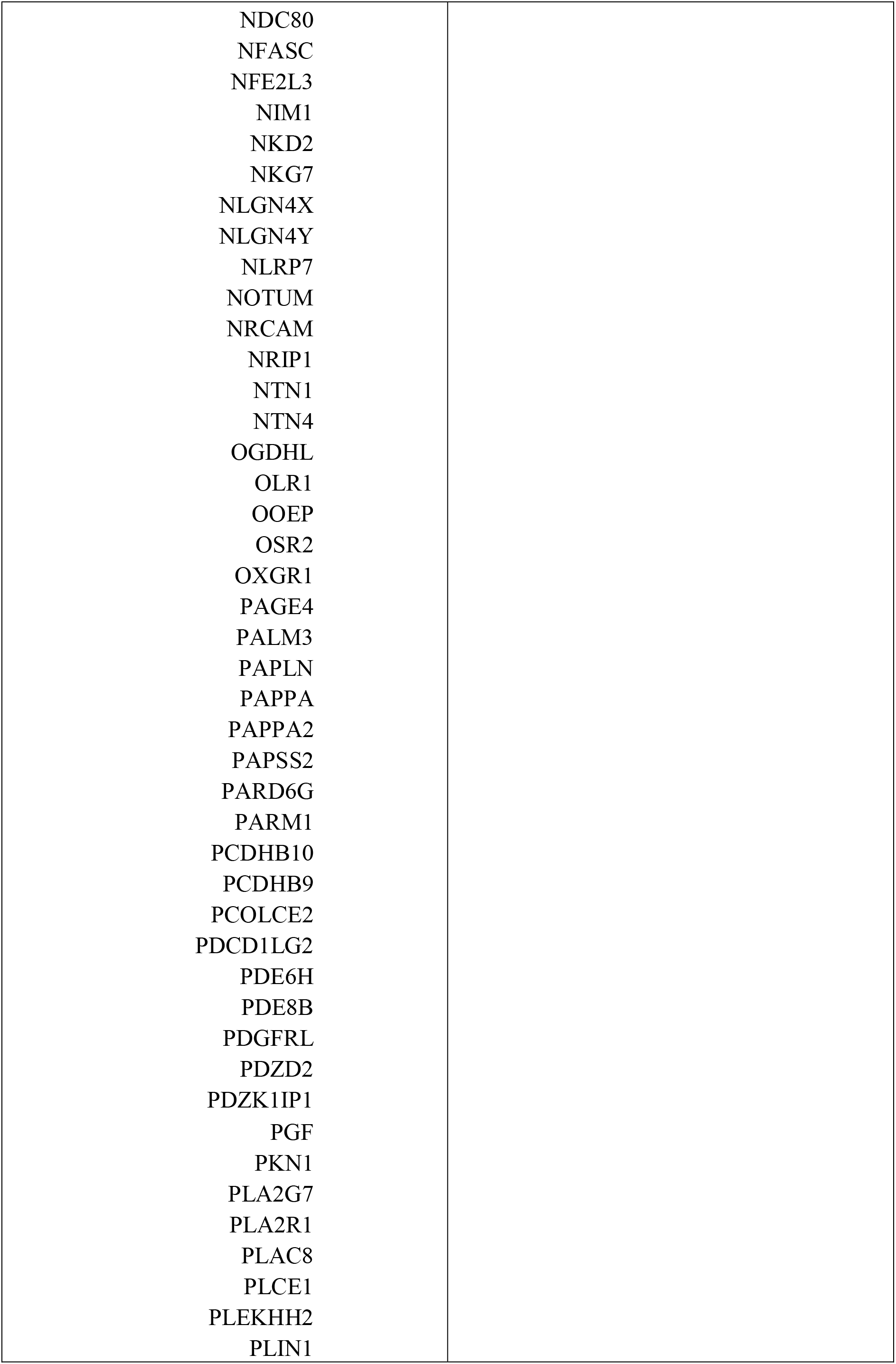
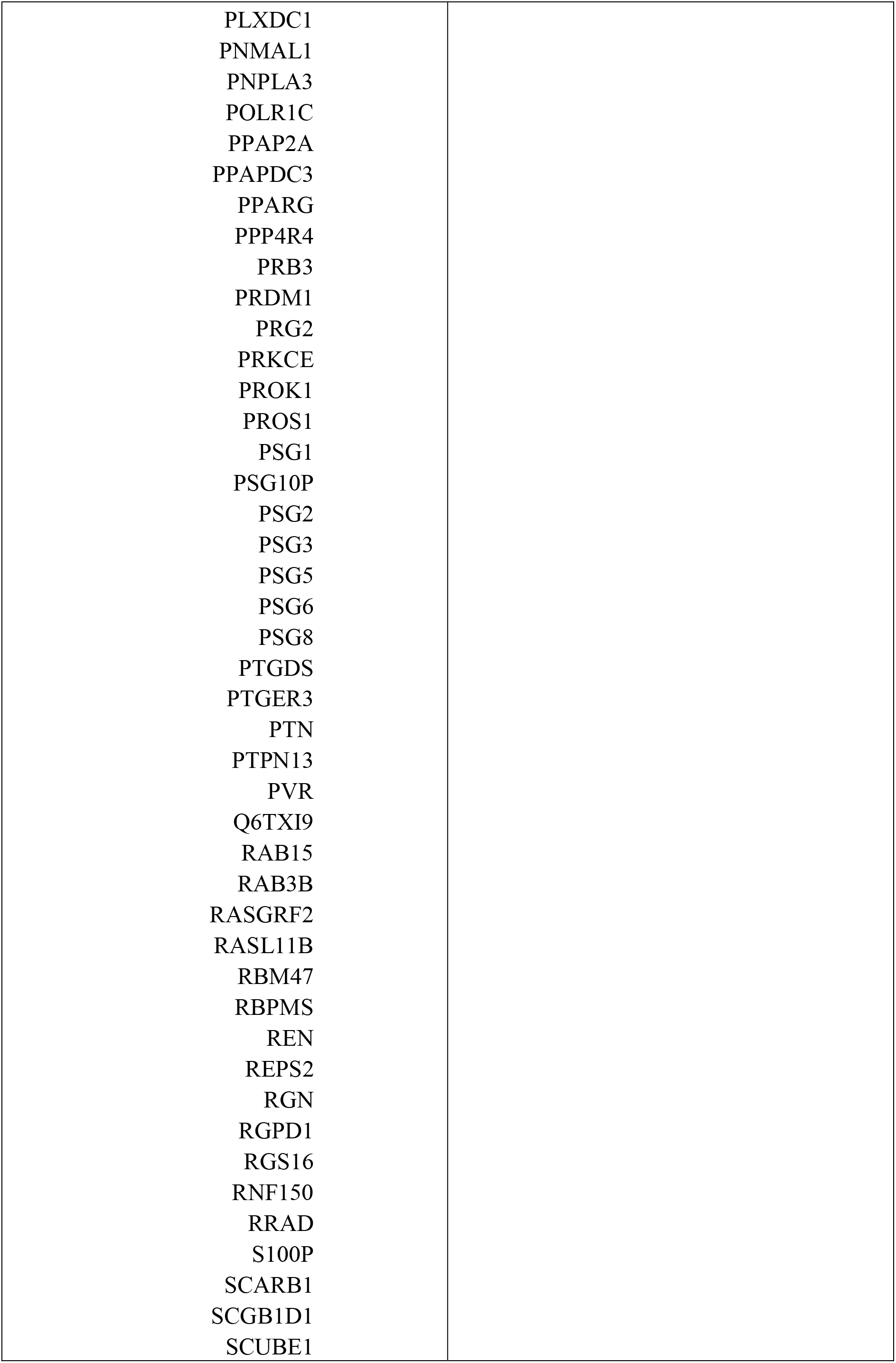
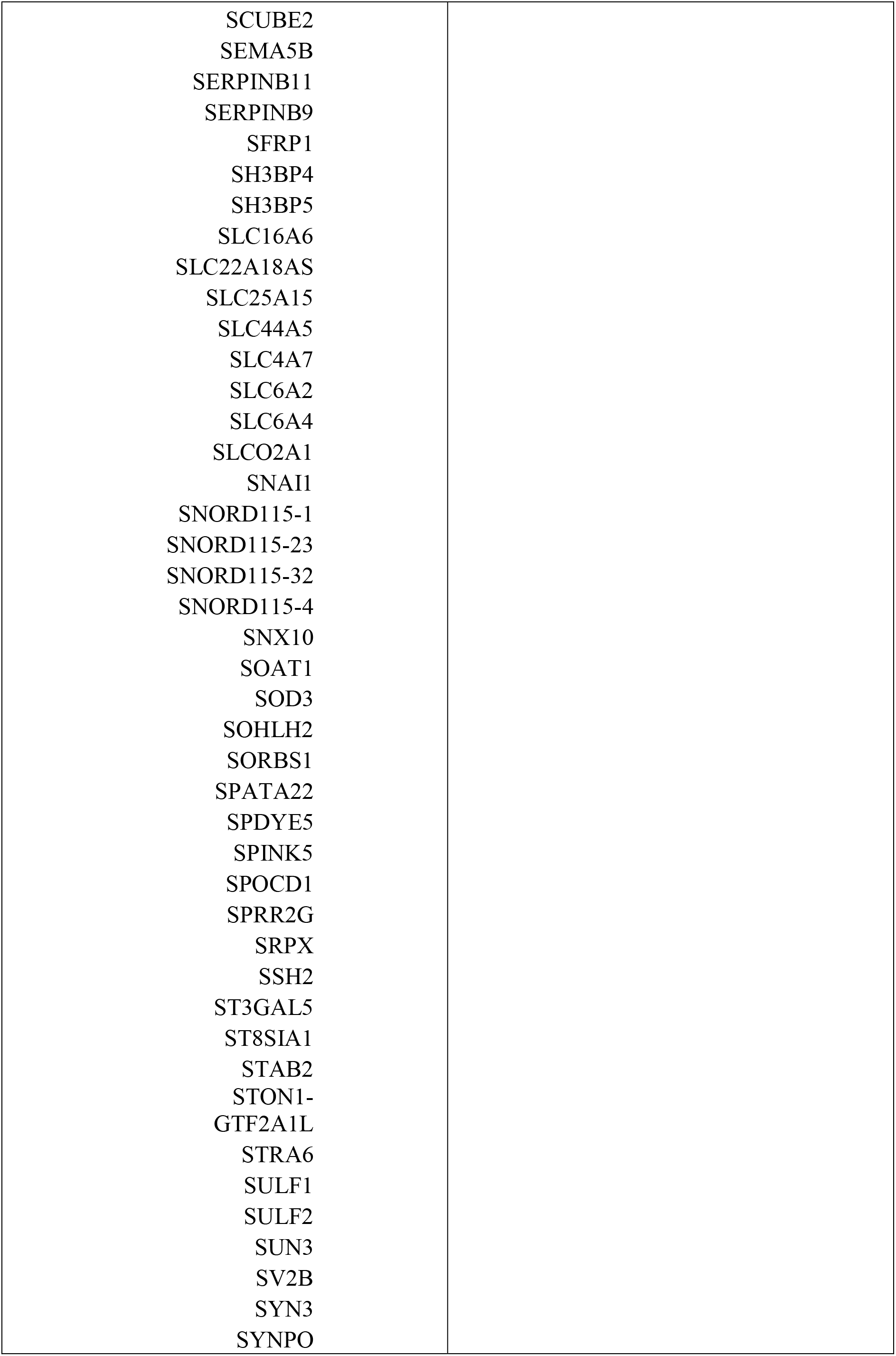
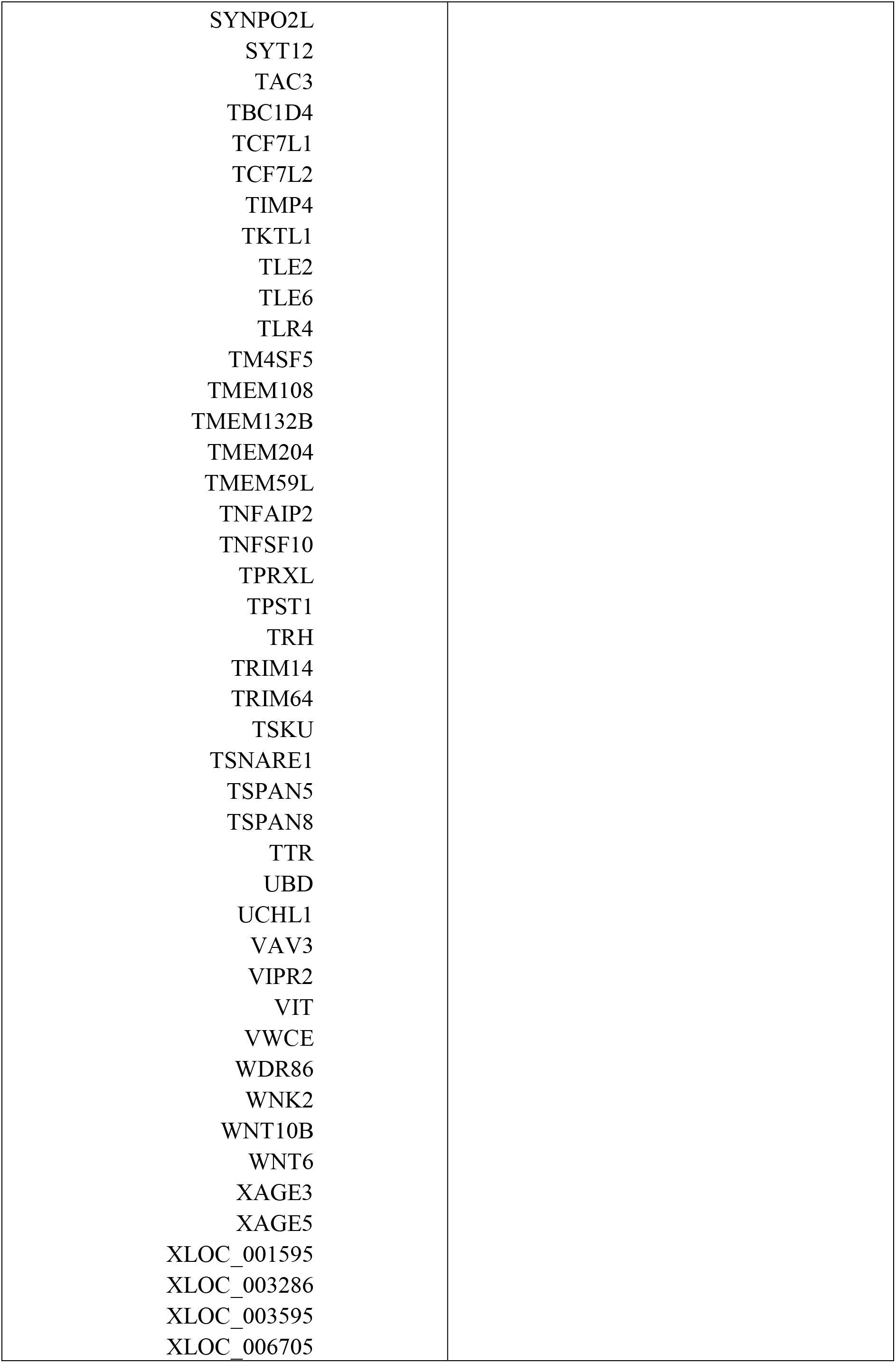
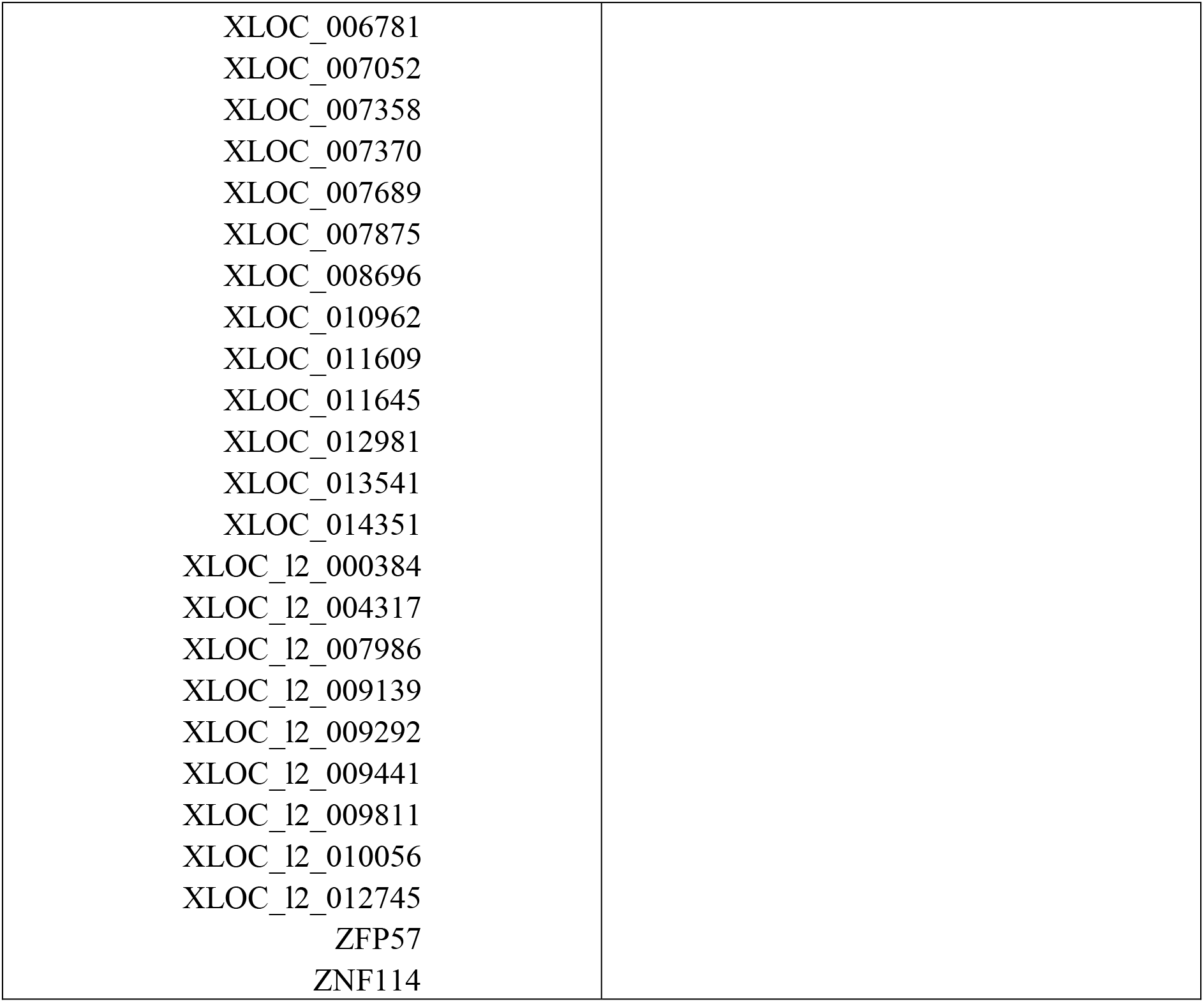
List of specific genes extracted from the ZAM zone after a transcriptomic analysis.

**Table S4:**
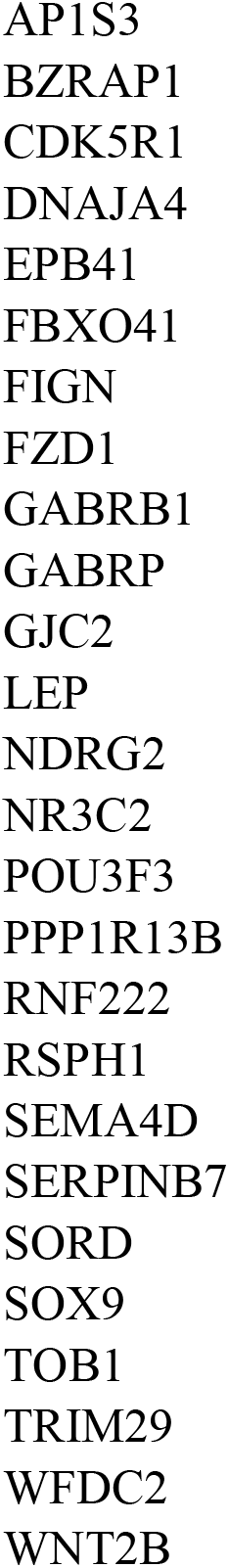
List of jointly hypermethylated genes in the ZAM choriodecidua and overexpressed in the ZAM amnion.

**Table S5:** List of disease mesh-terms associated with hypermethylated genes in the ZAM choriodecidua and overexpressed in the ZAM Amnion.

| MeSH-Term | MeSH-Term id(s) | p-value |
| --- | --- | --- |
| Disease Attributes | C23.550.291 | 8,17E-05 |
| Virilism | C23.888.971 | 1,52E-04 |
| Genetic Predisposition to Disease | C23.550.291.687.500 | 3,24E-04 |
| Disease Susceptibility | C23.550.291.687 | 4,59E-04 |
| Disease Progression | C23.550.291.656 | 4,84E-04 |
| Osteoporotic Fractures | C26.404.545 | 8,30E-04 |
| Endocrine Gland Neoplasms | C19.344, C04.588.322 | 8,85E-04 |
| Gliosis | C23.550.369 | 1,26E-03 |
| Testicular Diseases | C12.294.829 | 1,49E-03 |
| alpha-Thalassemia | C16.320.070.875.100, C15.378.071.141.150.875.100 | 1,50E-03 |
| Pancreatic Neoplasms | C19.344.421, C06.301.761, C06.689.667, C04.588.274.761, C04.588.322.475 | 1,50E-03 |
| Brain Neoplasms | C10.228.140.211, C10.551.240.250, C04.588.614.250.195 | 1,50E-03 |
| Anuria | C13.351.968.934.070, C13.351.968.419.078, C12.777.419.078, C12.777.934.141 | 1,65E-03 |
| Foot Deformities | C05.330 | 1,76E-03 |
| Lymphatic Metastasis | C23.550.727.650.560, C04.697.650.560 | 1,78E-03 |
| Pancreatic Diseases | C06.689 | 1,91E-03 |
| Skin Neoplasms | C17.800.882, C04.588.805 | 1,93E-03 |
| Neoplasms, Germ Cell and Embryonal | C04.557.465 | 2,06E-03 |
| Hyperandrogenism | C16.131.939.316.129.750, C12.706.316.064.500, C13.351.875.253.064.500, C19.391.119.129.750, C13.351.875.253.129.750, C16.131.939.316.064.500, C19.391.119.064.500, C12.706.316.129.750 | 2,14E-03 |
| Dementia | C10.228.140.380 | 2,16E-03 |
| Kwashiorkor | C18.654.521.500.708.626.505 | 2,30E-03 |
| Neurofibroma | C04.557.580.600.580 | 2,37E-03 |
| Cryptorchidism | C12.294.829.258, C16.131.939.258, C19.391.829.258, C12.706.258 | 2,46E-03 |
| Genital Neoplasms, Male | C12.758.409, C04.588.945.440, C12.294.260 | 2,55E-03 |
| Nerve Sheath Neoplasms | C04.557.580.600 | 2,62E-03 |
| Central Nervous System Neoplasms | C04.588.614.250, C10.551.240 | 2,68E-03 |
| Cardio-Renal Syndrome | C13.351.968.419.780.400, C14.280.434.156, C12.777.419.780.400 | 2,75E-03 |
| Sarcoma | C04.557.450.795 | 3,32E-03 |
| Urogenital Abnormalities | C16.131.939 | 3,38E-03 |
| Nervous System Neoplasms | C10.551 | 3,39E-03 |
| Nervous System Neoplasms | C04.588.614 | 3,42E-03 |
| Autoimmune Diseases | C20.111 | 3,44E-03 |
| Urogenital Abnormalities | C12.706 | 3,45E-03 |
| Mandibular Fractures | C26.404.750.467.441, C26.260.275.500.400.255 | 3,46E-03 |
| Mandibular Diseases | C05.500.607 | 3,62E-03 |
| Uremia | C13.351.968.419.936, C12.777.419.936 | 3,92E-03 |
| Jaw Fractures | C26.404.750.467, C26.260.275.500.400 | 4,01E-03 |
| Brain Infarction | C14.907.253.855.200, C10.228.140.300.775.200, C10.228.140.300.150.477, C14.907.253.092.477 | 4,03E-03 |
| Female Athlete Triad Syndrome | C05.116.198.579.304, C19.391.240 | 4,11E-03 |
| Mandibular Diseases | C07.320.610 | 4,14E-03 |
| Neoplasm Metastasis | C23.550.727.650, C04.697.650 | 4,35E-03 |
| Prostatic Neoplasms | C12.758.409.750, C12.294.260.750, C04.588.945.440.770, C12.294.565.625 | 4,49E-03 |
| Brain Diseases | C10.228.140 | 4,55E-03 |
| Sertoli Cell-Only Syndrome | C12.294.365.700.754 | 4,60E-03 |
| Pathologic Processes | C23.550 | 4,63E-03 |
| Osteophyte | C05.116.540.310.800 | 4,72E-03 |
| Temporomandibular Joint Disorders | C05.500.607.221.897, C07.320.610.291.897, C05.550.905, C05.651.243.897, C07.678 | 4,75E-03 |
| Pleural Effusion | C08.528.652 | 4,76E-03 |
| Adrenocortical Adenoma | C19.344.078.265.500, C19.053.347.500.500, C19.053.098.265.500, C04.588.322.078.265.500 | 4,80E-03 |
| Pseudohypoaldosteronism | C16.320.565.861.770, C13.351.968.419.815.770, C12.777.419.815.770, C18.452.648.861.770 | 4,84E-03 |
| Premenstrual Syndrome | C23.550.568.968 | 4,84E-03 |
| 46, XX Disorders of Sex Development | C12.706.316.064, C19.391.119.064, C13.351.875.253.064, C16.131.939.316.064 | 4,85E-03 |
| Cranio-mandibular Disorders | C05.500.607.221, C07.320.610.291, C05.651.243 | 4,96E-03 |
| Osteoporosis | C05.116.198.579 | 5,04E-03 |
| Prostatic Diseases | C12.294.565 | 5,29E-03 |
| alpha-Thalassemia | C16.320.365.826.100, C15.378.420.826.100 | 5,29E-03 |
| Neurodegenerative Diseases | C10.574 | 5,32E-03 |
| Tracheomalacia | C17.300.182.895.500, C16.131.621.953.500, C08.907.796.500, C05.182.895.500 | 5,48E-03 |
| Tracheobronchomalacia | C08.907.796 | 5,48E-03 |
| Anhedonia | C23.888.592.604.039, C10.597.606.057 | 5,61E-03 |
| Heart Septal Defects, Ventricular | C16.131.240.400.560.540, C14.280.400.560.540, C14.240.400.560.540 | 5,87E-03 |
| Neurilemmoma | C04.557.580.625.650.595 | 6,08E-03 |
| Genital Diseases, Male | C12.294 | 6,11E-03 |
| Infertility, Male | C12.294.365.700 | 6,15E-03 |
| Pseudotumor Cerebri | C10.228.140.631.750 | 6,16E-03 |
| Endocrine System Diseases | C19 | 6,43E-03 |
| Brain Ischemia | C14.907.253.092, C10.228.140.300.150 | 6,44E-03 |
| Tooth Diseases | C07.793 | 6,50E-03 |
| Respiratory Tract Diseases | C08 | 6,65E-03 |
| Cholestasis | C06.130.120.135 | 6,80E-03 |
| Kidney Failure, Chronic | C13.351.968.419.780.750.500, C12.777.419.780.750.500 | 6,82E-03 |
| Tracheobronchomalacia | C16.131.621.953, C17.300.182.895, C05.182.895 | 6,84E-03 |
| Urinary Tract Infections | C01.539.895, C13.351.968.892, C12.777.892 | 6,85E-03 |
| Maxillofacial Injuries | C26.260.275.500 | 7,02E-03 |
| Autoimmune Diseases of the Nervous System | C20.111.258, C10.114 | 7,16E-03 |
| Encephalomyelitis, Autoimmune, Experimental | C20.111.258.625.300, C10.114.703.300, C10.314.350.250, C10.228.140.695.562.250 | 7,19E-03 |
| Immune System Diseases | C20 | 7,22E-03 |
| Ossification of Posterior Longitudinal Ligament | C05.116.900.480, C23.550.751.500 | 7,31E-03 |
| Cerebrovascular Disorders | C10.228.140.300, C14.907.253 | 7,69E-03 |
| Osteochondritis | C17.300.182.520, C05.182.520 | 7,77E-03 |
| Chondrosarcoma | C04.557.450.795.300, C04.557.450.565.280 | 7,78E-03 |
| Pelvic Inflammatory Disease | C01.539.635.500 | 8,08E-03 |
| Neurilemmoma | C04.557.580.600.610.595, C04.557.465.625.650.595 | 8,20E-03 |
| Refeeding Syndrome | C18.654.521.687 | 8,21E-03 |
| Acanthosis Nigricans | C17.800.621.430.530.100 | 8,24E-03 |
| Urogenital Neoplasms | C12.758 | 8,25E-03 |
| Carcinoma, Basal Cell | C04.557.470.565.165, C04.557.470.200.165 | 8,36E-03 |
| Nephritis, Interstitial | C12.777.419.570.643, C13.351.968.419.570.643 | 8,50E-03 |
| Nervous System Autoimmune Disease, Experimental | C10.114.703, C20.111.258.625 | 8,55E-03 |
| Teratocarcinoma | C04.557.465.900 | 8,59E-03 |
| Hyperphosphatemia | C18.452.750.199 | 8,72E-03 |
| Pelvic Infection | C01.539.635 | 8,72E-03 |
| Pleural Diseases | C08.528 | 8,98E-03 |
| Demyelinating Autoimmune Diseases, CNS | C10.228.140.695.562 | 9,11E-03 |
| Neuroendocrine Tumors | C04.557.580.625.650 | 9,17E-03 |
| Central Nervous System Diseases | C10.228 | 9,19E-03 |
| Hereditary Autoinflammatory Diseases | C17.800.827.368 | 9,39E-03 |
| Neuroendocrine Tumors | C04.557.465.625.650 | 9,42E-03 |
| Heart Failure, Systolic | C14.280.434.676 | 9,56E-03 |
| TDP-43 Proteinopathies | C10.574.950, C18.452.845.800 | 9,75E-03 |
| Demyelinating Autoimmune Diseases, CNS | C10.314.350 | 9,83E-03 |
| Carcinogenesis | C04.697.098, C23.550.727.098 | 9,88E-03 |
| Chromosome Inversion | C23.550.210.190 | 9,88E-03 |
| Esophageal and Gastric Varices | C06.552.494.414, C06.405.117.240 | 9,90E-03 |
| Protozoan Infections, Animal | C03.752.625, C03.701.688, C22.674.710 | 9,94E-03 |

**Table S6:** List of jointly hypermethylated genes in the ZAM amnion and overexpressed in the ZAM choriodecidua.

|  |  |  |
| --- | --- | --- |
| ADAMTS1 | HPCAL1 | OOEP |
| ADAMTS5 | HPGD | PAGE4 |
| ANTXR1 | HSD3B1 | PAPPA2 |
| APOA1 | ICAM2 | PAPSS2 |
| AR | IL15 | PARD6G |
| ATP13A4 | IL1RL1 | PDCD1LG2 |
| BIN2 | IL20RB | PLEKHH2 |
| C1orf130 | ISM2 | PPAP2A |
| C1QTNF1 | ITGA1 | PPAPDC3 |
| C2CD4B | JAM2 | PPARG |
| C7orf58 | JAZF1 | PPP4R4 |
| CDA | KCNJ16 | PRB3 |
| CEACAM3 | KIAA1324 | PRG2 |
| COL14A1 | KITLG | PROK1 |
| CORO2A | KLK3 | PTGER3 |
| CXCR6 | LAIR2 | SCARB1 |
| CYP11A1 | LGALS14 | SERPINB11 |
| DENND2A | LIFR | SLC44A5 |
| DIO2 | LINGO2 | SLC6A2 |
| DSCR8 | LIPG | SOAT1 |
| EEPD1 | MAGEA8 | SOHLH2 |
| ENTPD8 | MAPT | SSH2 |
| FAM153B | MCAM | STAB2 |
| FHOD3 | MFSD2A | SV2B |
| FIGNL2 | MMP11 | TLR4 |
| FLJ43663 | MMP7 | TM4SF5 |
| GCM1 | NFASC | TMEM132B |
| GDF15 | NFE2L3 | TNFSF10 |
| GH1 | NKD2 | TPST1 |
| GKN1 | NKG7 | TRIM14 |
| GPC4 | NLGN4X | TSKU |
| GTSF1 | NLRP7 | TSNARE1 |
| GYLTL1B | OGDHL | TSPAN8 |
| HMGB3 | OLR1 | VAV3 |
| HN1 |  | XAGE3 |
|  |  | ZFP57 |

**Table S7:**
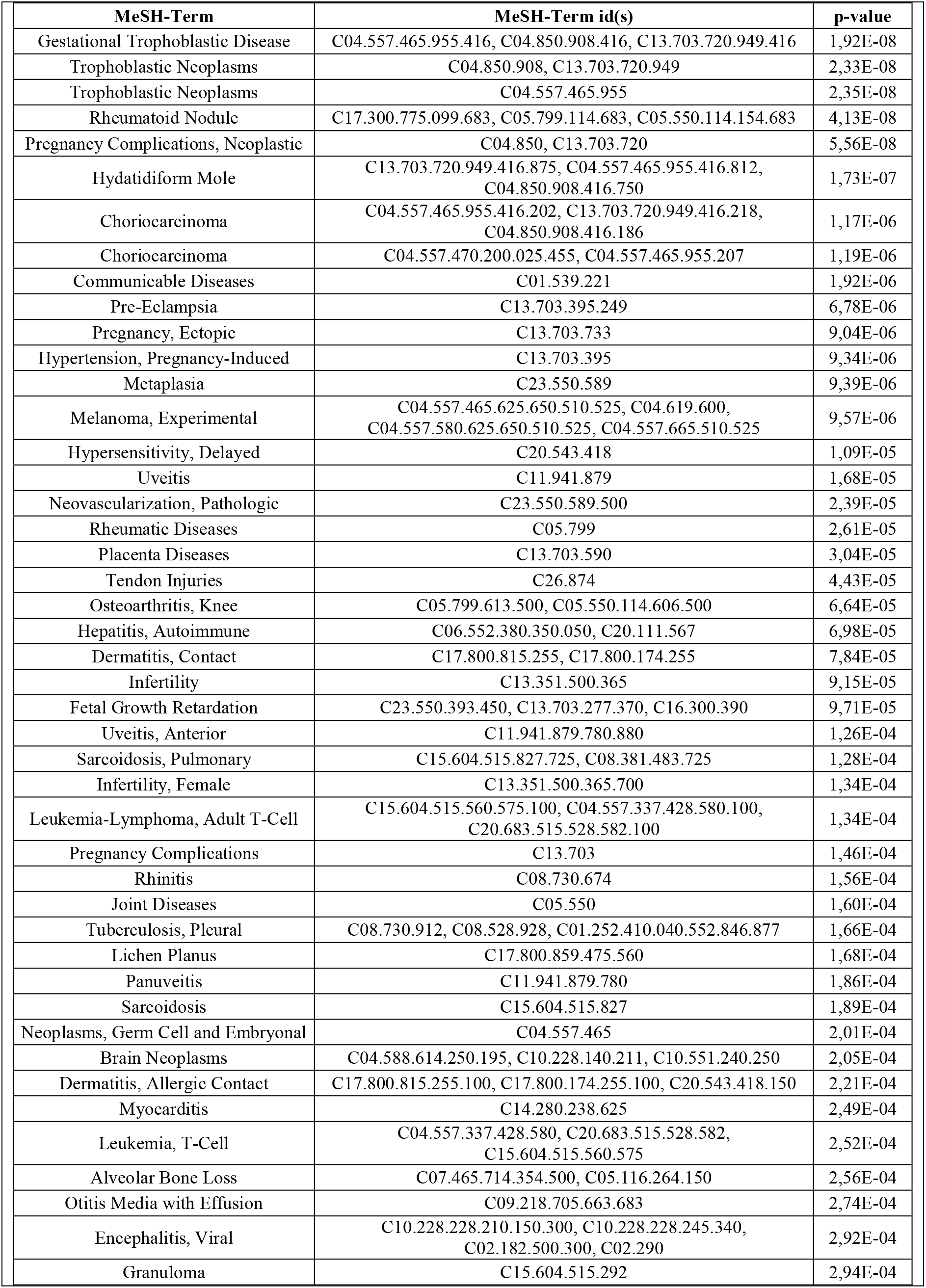
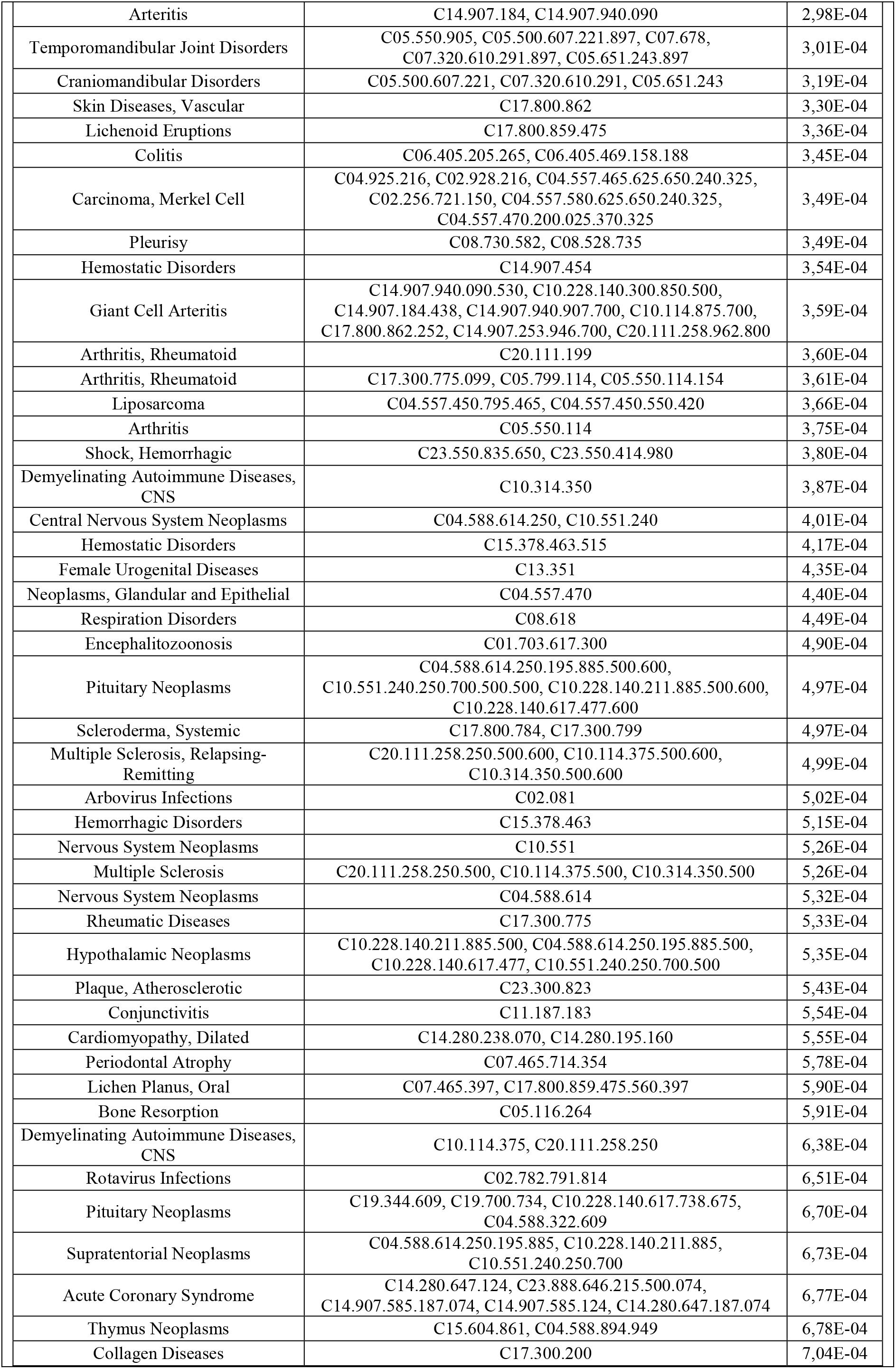
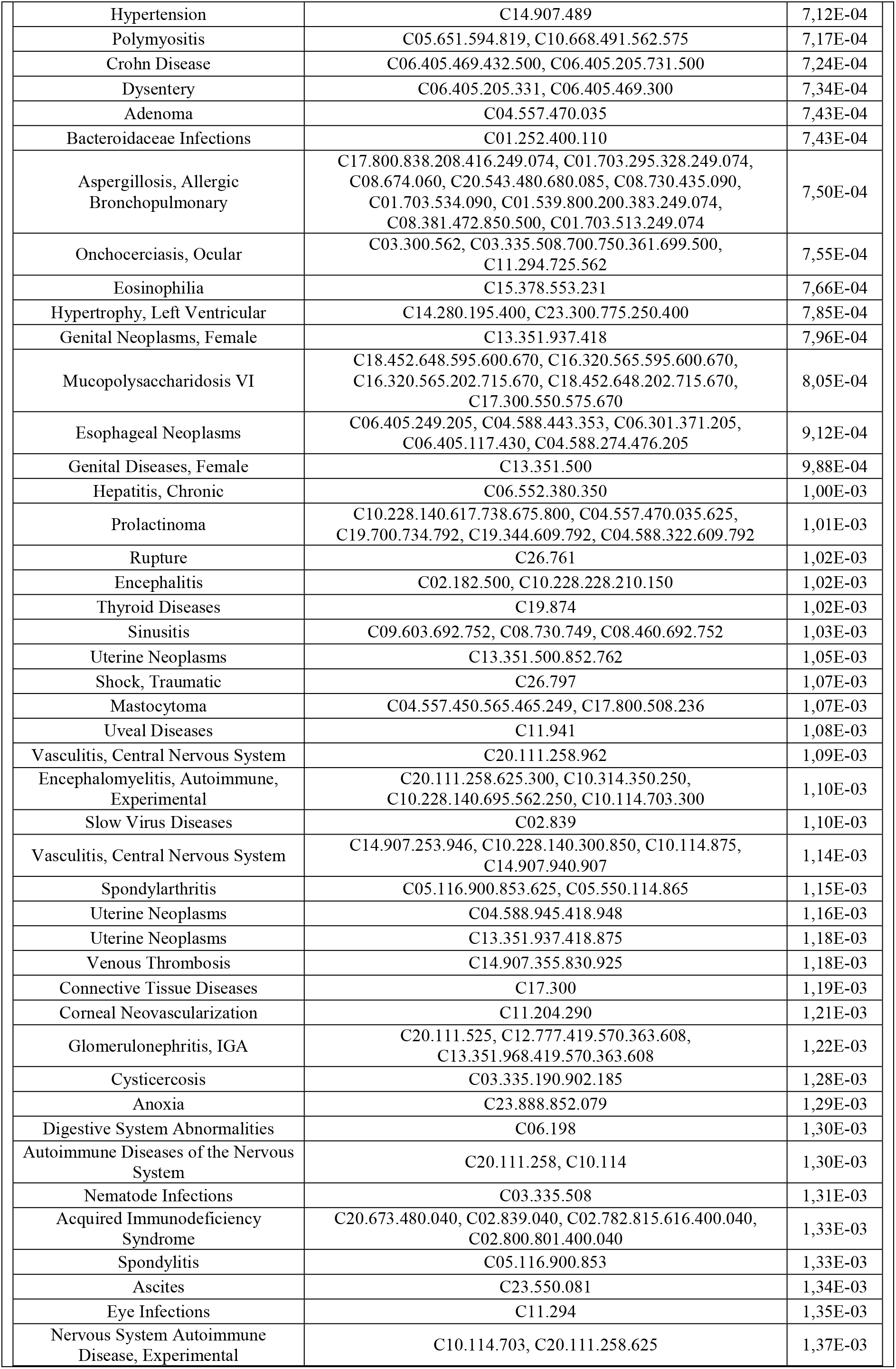
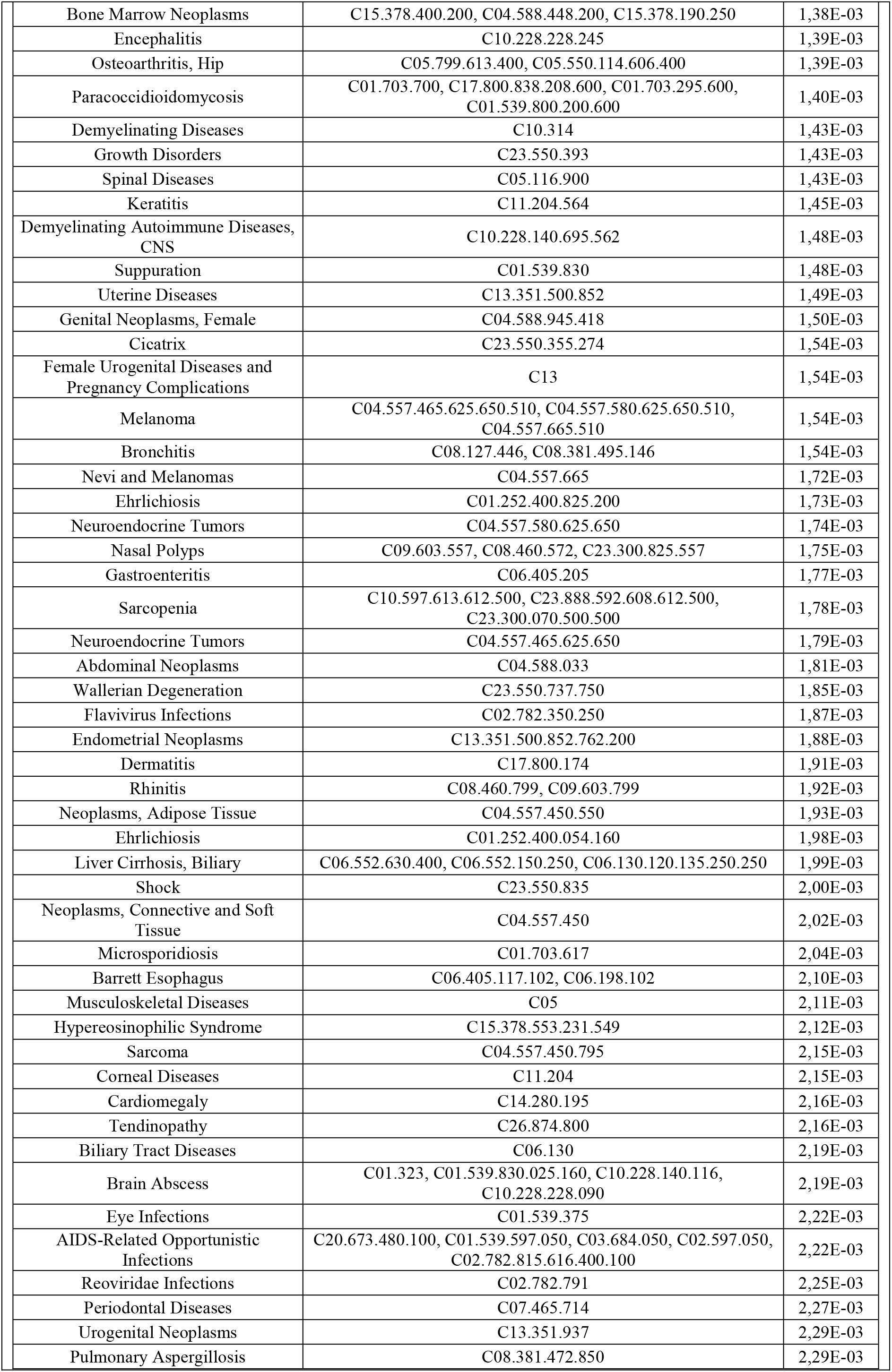
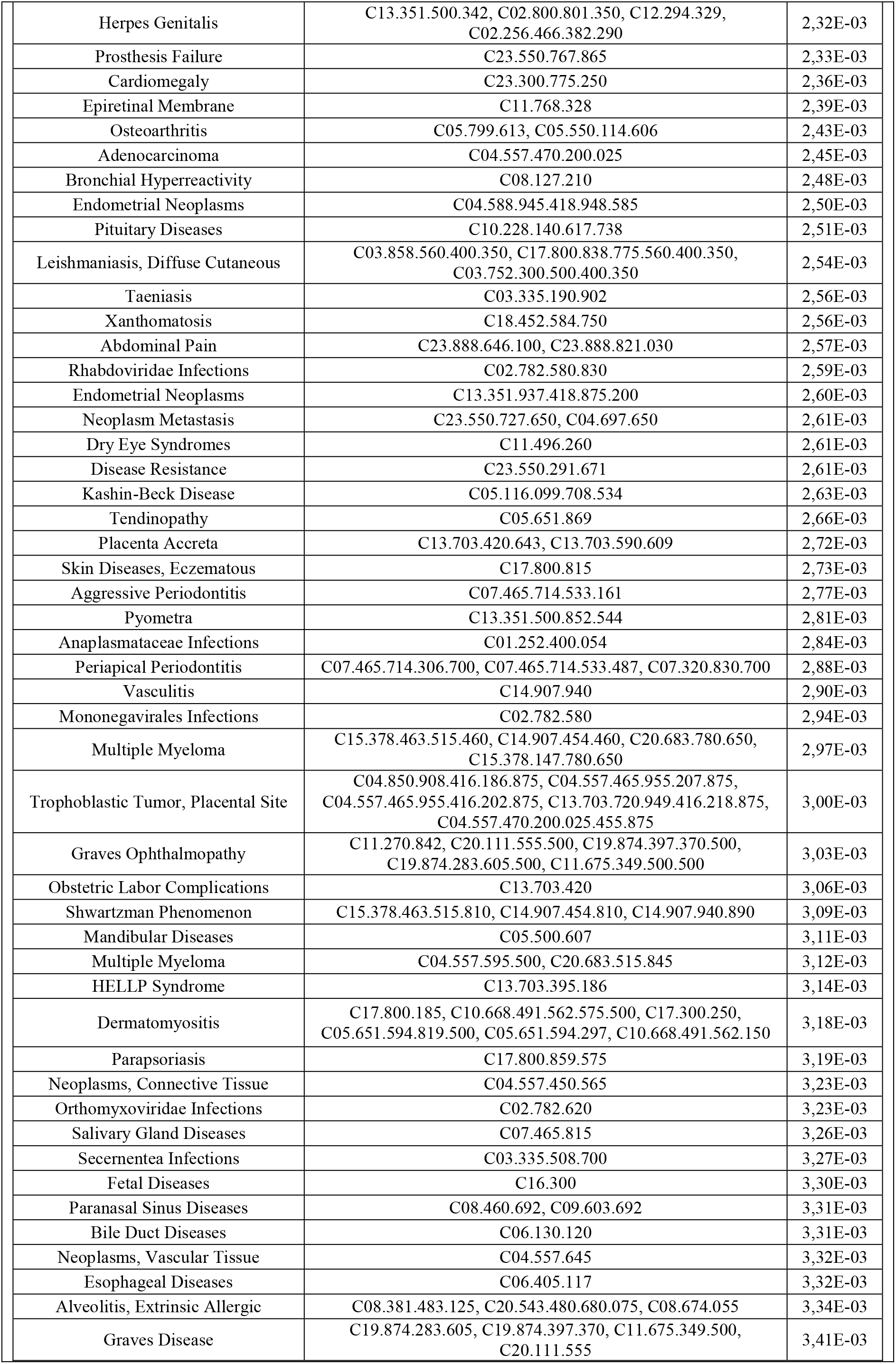
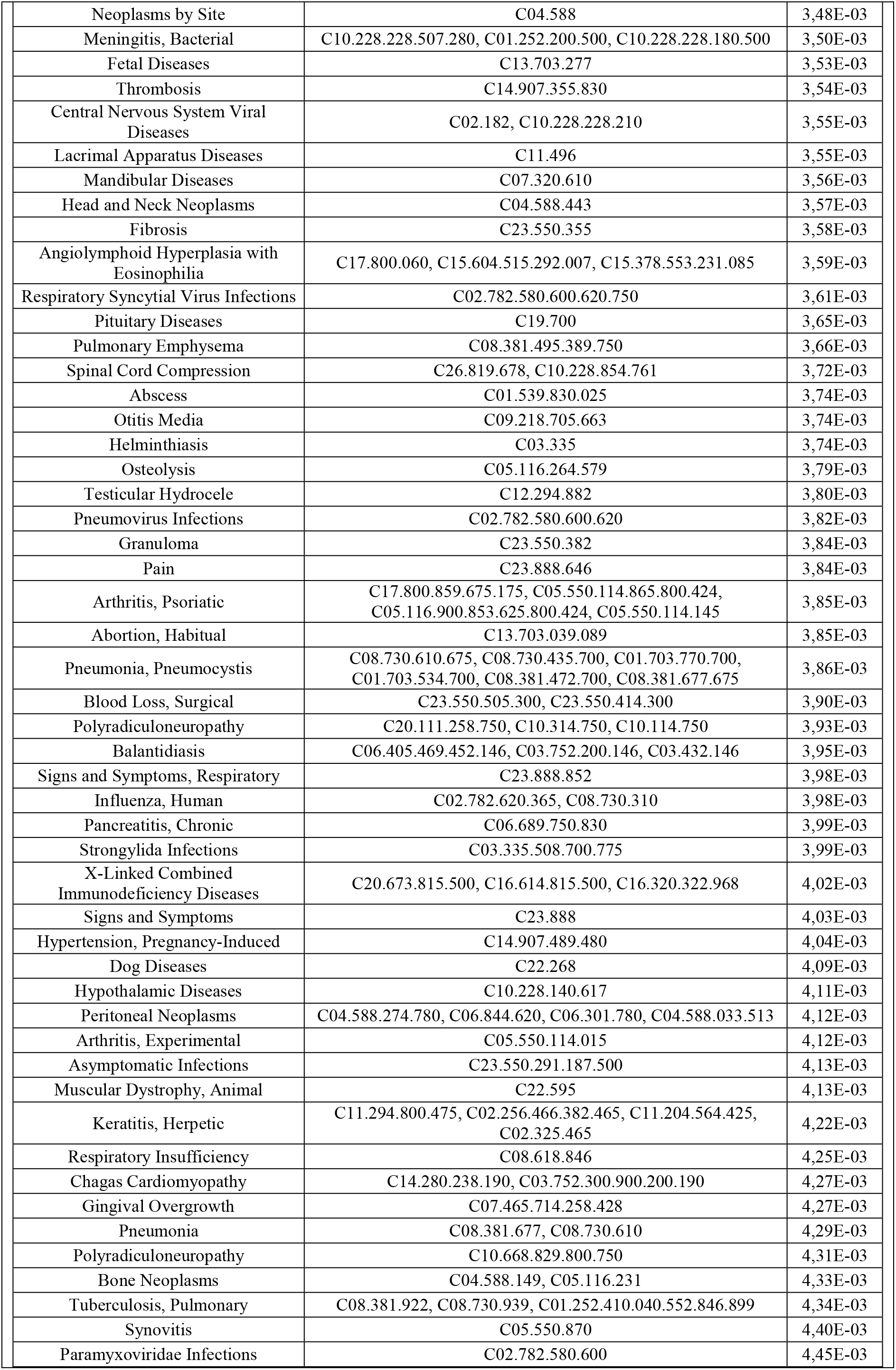
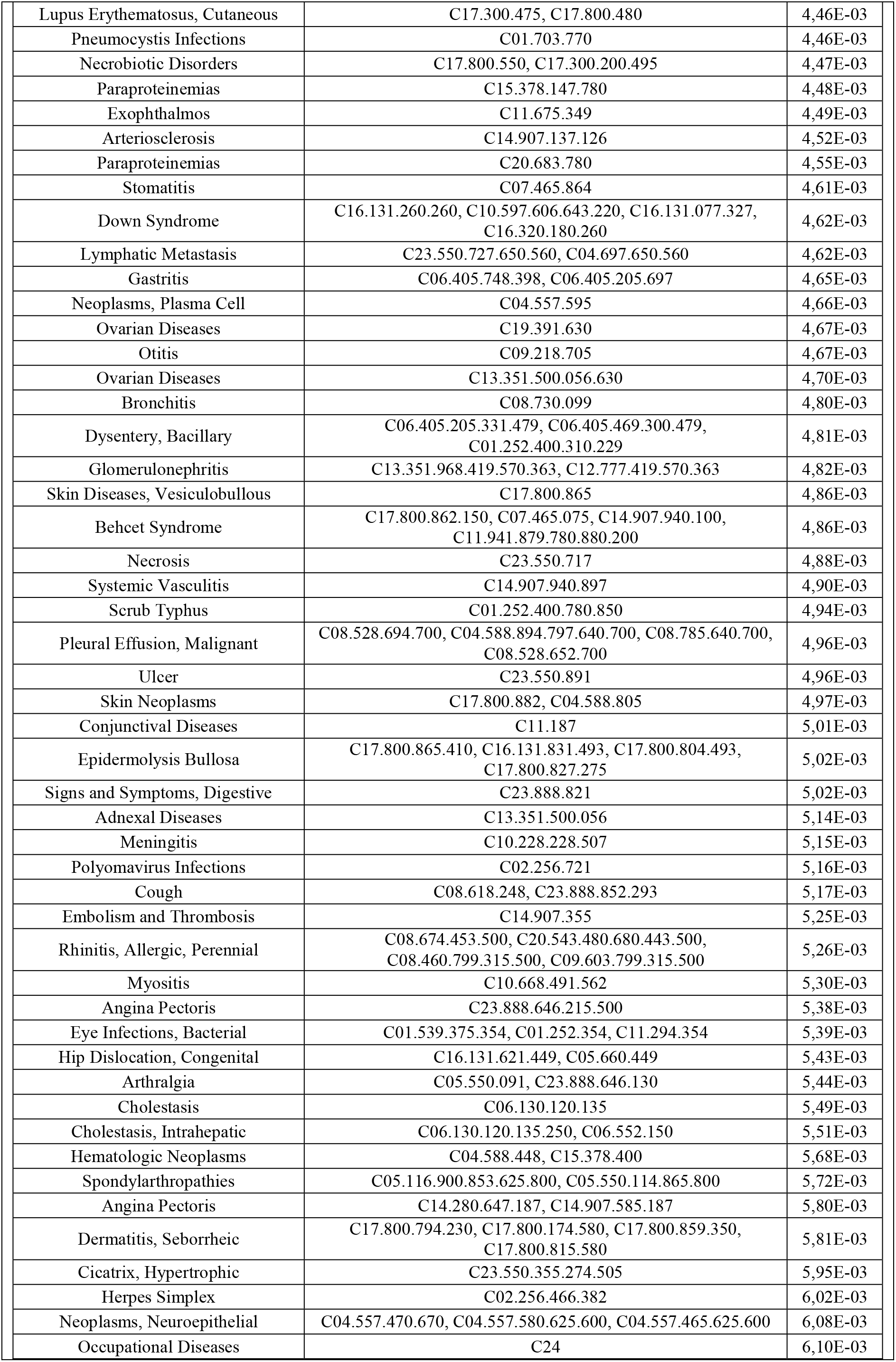
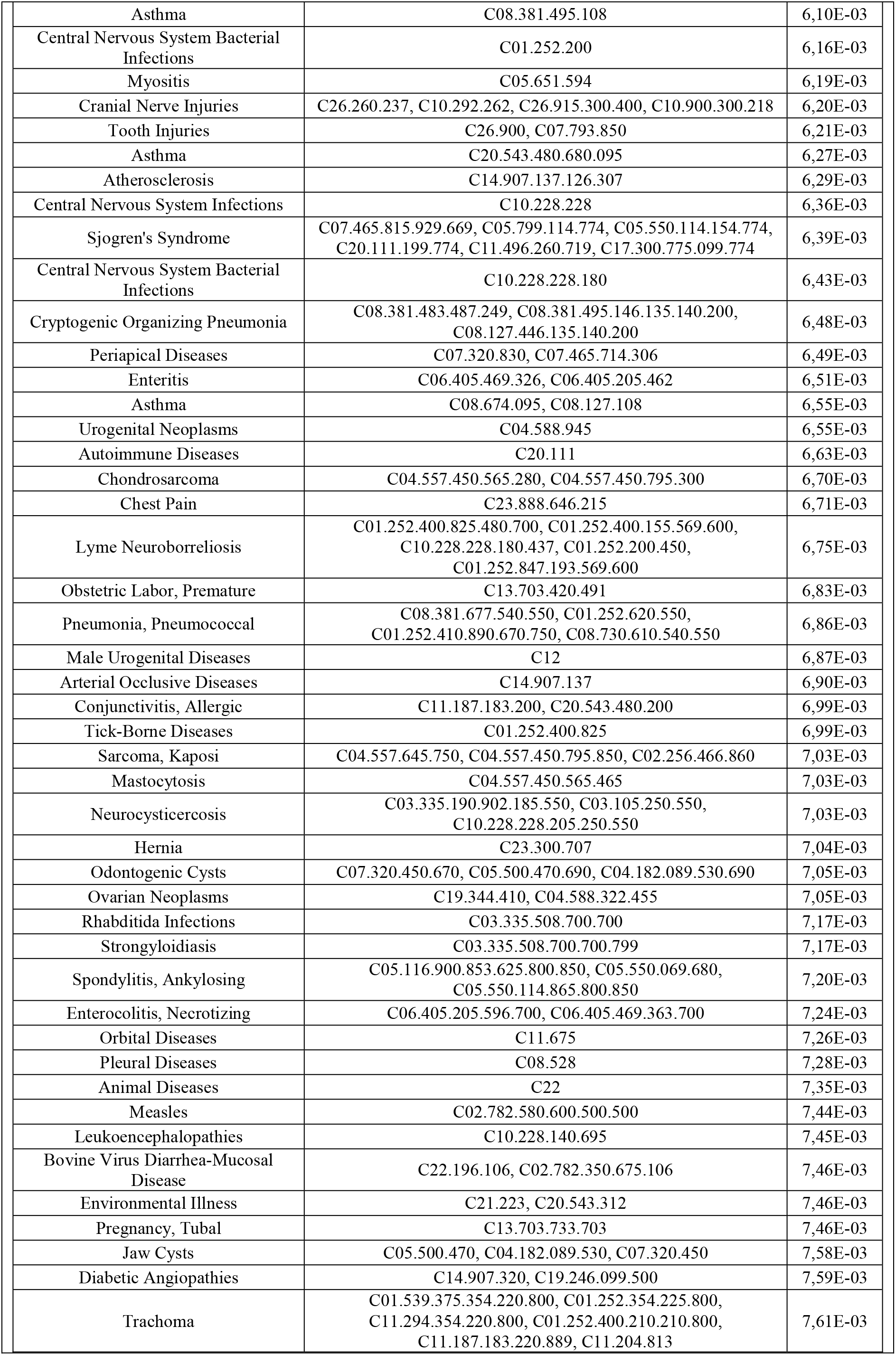
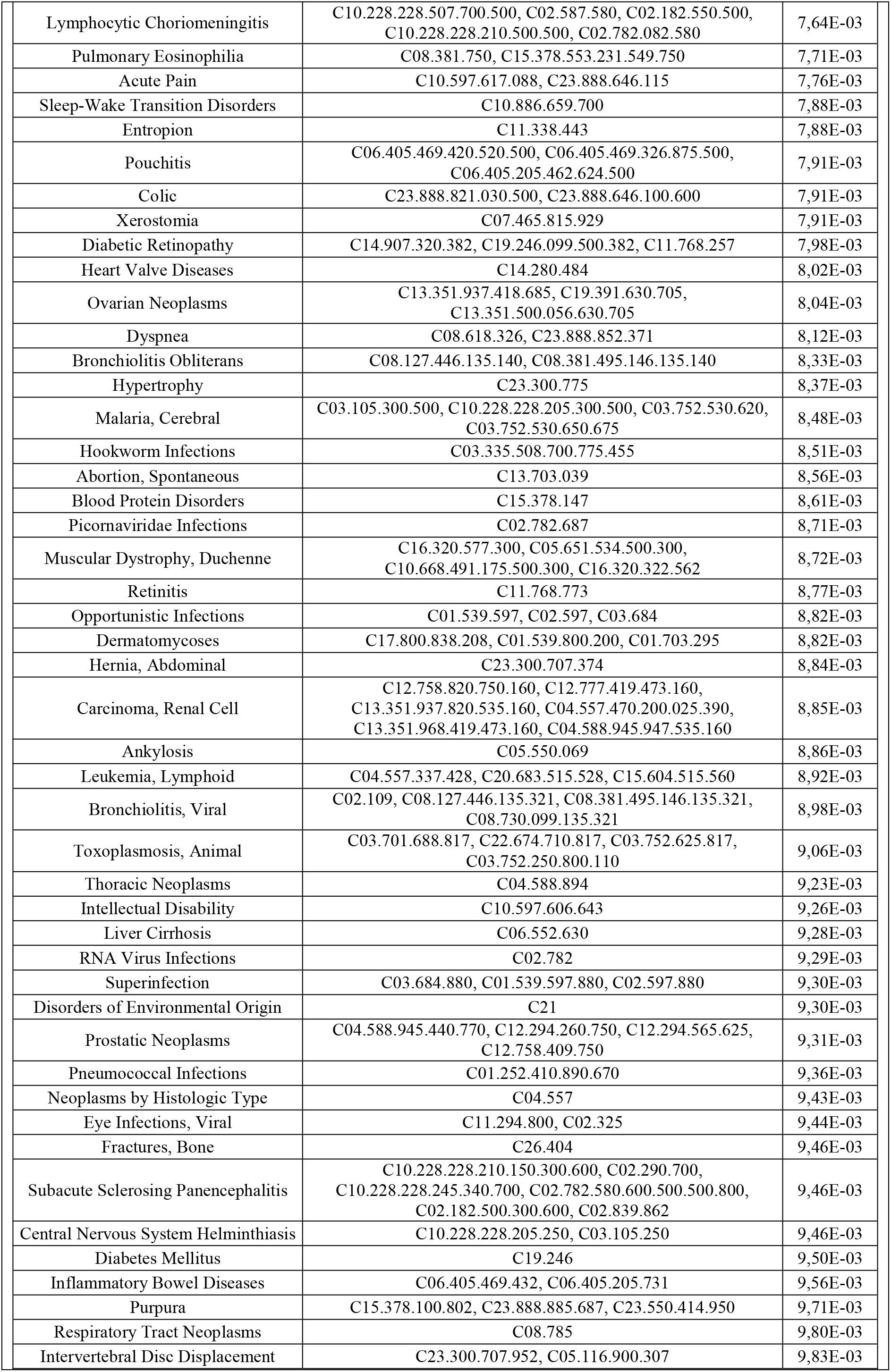
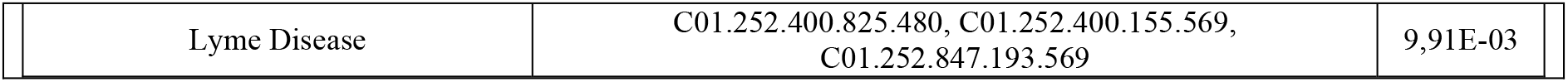
List of disease mesh-terms associated with hypermethylated genes in the ZAM amnion and overexpressed in the ZAM choriodecidua.

